# A painless nerve growth factor variant uncouples nociceptive and neurotrophic TrkA signaling

**DOI:** 10.1101/2025.11.18.686377

**Authors:** Shailesh Kumar, Harry A. Jung, Gerard Walker, Jaimin K. Rana, Rui Fu, Minh Triết Hồng, Francesca Malerba, Jyotsna Kumar, Vikas Navratna, Wonhyeuk Jung, Sean J. Miller, Lukas Kruckenhauser, Simona Capsoni, Brian P. Hafler, Antonino Cattaneo, Yamuna Krishnan, Shyamal Mosalaganti, Moitrayee Bhattacharyya

## Abstract

Nerve growth factor (NGF) binding to the receptor tyrosine kinase, TrkA, drives neurotrophic signaling essential for neuronal development and survival. This interaction simultaneously drives peripheral pain, making this pathway an attractive but complicated therapeutic target for chronic pain. By integrating single-molecule microscopy, structural and electrophysiology analyses, with a human NGF variant, NGF^painless^, which retains neurotrophic effects but abolishes pain, we delineate the molecular mechanisms that bias TrkA signaling towards neurotrophic functions without triggering nociception. We show that, unlike wild-type NGF, NGF^painless^ fails to sensitize TRPV1 channels to capsaicin, thus disengaging TrkA from the nociceptive pathway. We further show that this selective loss of nociceptive TrkA signaling by NGF^painless^ results from its reduced ability to activate PLCγ1 and trigger calcium release compared to NGF, while still preserving the ERK and AKT signaling essential for neurotrophic functions. This biased signaling arises from reduced electrostatic complementarity at the TrkA:NGF^painless^ complex interface, which shortens the lifetime of this functional complex on native membranes. Mutations in TrkA that restore the electrostatic complementarity at the TrkA:NGF^painless^ interface eliminate biased signaling. This mechanistic understanding of TrkA binding by NGF^painless^, and how it differs from NGF, will spur the development of two therapeutic classes of molecules - one that selectively suppresses nociceptive signaling while preserving neurotrophic functions in chronic pain, and another that enhances neurotrophic activity without evoking peripheral pain in neurodegenerative conditions.

## Introduction

About one in ten people who are prescribed opioids for chronic pain end up with dependence and addiction^1–4^. This underscores the urgent need to delineate the molecular mechanisms of pathways underlying chronic pain, enabling the development of non-addictive pain medication. In this regard, the receptor tyrosine kinase TrkA and its ligand, nerve growth factor (NGF), are valuable therapeutic targets^5,6^. TrkA activation by NGF drives the beneficial neurotrophic signaling that promotes neuronal growth and survival, yet it simultaneously triggers peripheral pain by activating nociceptive pathways^6–8^. While inhibiting TrkA:NGF signaling with antagonists relieves pain in diverse pathologies^9–18^, these inhibitors and antibodies have shown adverse effects in clinical trials^8,11,13,18^, as they unavoidably block the beneficial neurotrophic functions. Therefore, identifying molecular mechanisms that uncouple the nociceptive from the neurotrophic outcomes of TrkA signaling is key to successfully targeting the TrkA:NGF pathway therapeutically.

Humans with a rare mutation in NGF, i.e., R100W, in hereditary sensory and autonomic neuropathy type V (HSANV), exhibit loss of pain perception yet show no obvious neurological defects^19–25^. This led to the design of an NGF variant bearing R100E+P61S mutations, denoted NGF^painless^, which is easier to produce in high quality and yield compared to the naturally occurring NGF^R100W,19,24,26–31^. NGF^painless^ shows comparable stability to NGF (**Supplementary Fig. 1**), and recapitulates the phenotype of NGF^R100W^, as it does not trigger pain or hyperalgesia in animal models and clinical trials^19,24,27–30,38–40^. Unlike TrkA mutations that diminish pain perception and cause neurological abnormalities^21,32–37^, the “painless” variant selectively abolishes nociception while preserving neurotrophic functions^19,24,26–30^. However, the structural and mechanistic basis for how NGF^painless^ selectively permits TrkA to recruit neurotrophic, but not nociceptive signaling pathways remain unresolved.

TrkA consists of an extracellular domain (ECD), which is connected to an intracellular kinase domain (KD) by a single-pass transmembrane domain (TM) and extra-/intra-cellular juxtamembrane regions (EJM/IJM)^5,6^. The ECD contains two immunoglobulin-like regions (Ig1 and Ig2), with Ig2 forming the NGF-binding interface^41,42^. NGF binding promotes TrkA oligomerization and activation, leading to autophosphorylation at two tyrosine residues, Y490 and Y785, on the receptor. Activated TrkA then phosphorylates and activates three downstream signaling pathways, namely, Ras–MAPK/ERK, PI3K/AKT, and PLCγ1 (**Fig. 1a**). When NGF activates TrkA, it also drives nociception by sensitizing ion channels, such as TRPV1, to capsaicin and other noxious stimuli^43–45^. This process has been shown to be critically mediated by PLCγ1 activation^44,46–48^.

**Figure 1:**
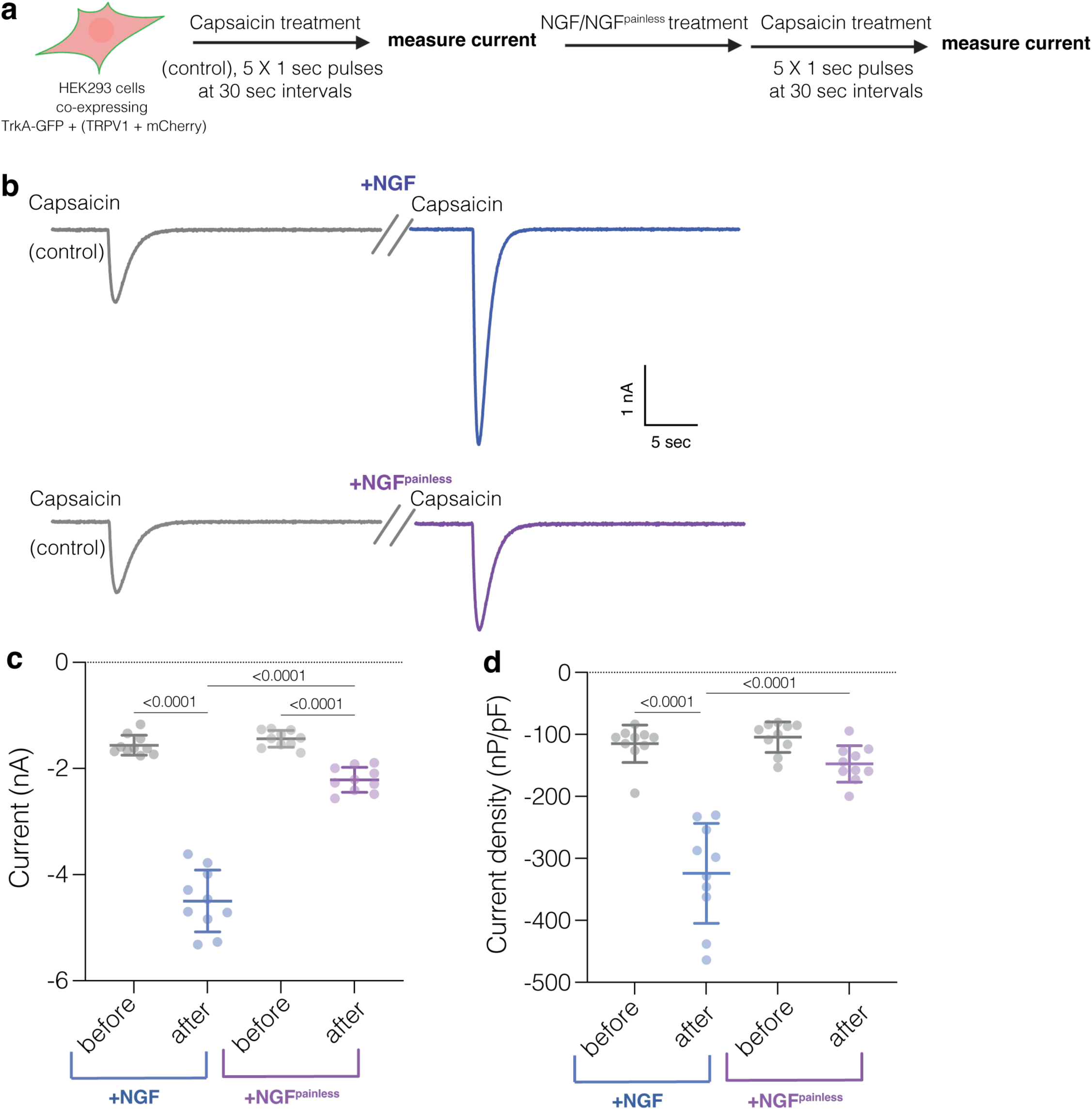
Capsaicin-evoked currents in TRPV1 upon treatment with NGF and NGF^painless^. **a.** Schematic showing the experimental design for electrophysiological recordings. **b.** Representative capsaicin-evoked inward currents (1 sec pulses of 10 μM capsaicin) in a voltage-clamped HEK293 cell co-expressing TrkA and TRPV1, before and after treatment with 100 ng/ml of NGF (top panel) or NGF^painless^ (bottom panel) for 15 min at 25 °C. **c.** Comparison of capsaicin-evoked currents from N=10 cells before (gray) and after treatments with NGF (blue) or NGF^painless^ (purple, -60 mV holding potential; also **Supplementary** Fig. 2). Experimental conditions are the same as in (b). **d.** Comparison of capsaicin-evoked current densities (defined as whole-cell membrane current divided by cell membrane capacitance) from the same N=10 cells in c, before and after treatments with NGF or NGF^painless^ (-60 mV holding potential). Experimental conditions are the same as in (b-c).

Here, using *in vitro* and live cell single-molecule microscopy, electrophysiology, biochemistry, and single-particle cryogenic electron microscopy (cryo-EM), we show that unlike NGF, NGF^painless^ biases TrkA signaling to selectively block nociceptive pathways. We demonstrate that, unlike NGF, NGF^painless^ fails to sensitize TRPV1 channels to noxious stimuli such as capsaicin. We further show that this selective loss of nociceptive TrkA signaling by NGF^painless^ results from its reduced ability to activate PLCγ1 and subsequent calcium release from intracellular stores, while maintaining ERK and AKT activation essential for the neurotrophic functions. We uncovered that the TrkA:NGF^painless^ complex exhibits altered kinetics on native membranes, with lower dimer/multimer lifetimes, compared to TrkA:NGF. Finally, by providing high-resolution structures of TrkA:NGF and TrkA:NGF^painless^ complexes, we reveal the structural rationale for the reduced lifetime of the TrkA:NGF^painless^ complex, which biases downstream signaling. We further validated this structural mechanism by introducing structure-guided complementary mutations in TrkA, which no longer exhibited biased signaling by NGF^painless^.

Taken together, our results define how one might ligand TrkA to selectively inhibit nociception yet preserve neurotrophic outcomes or enhance neurotrophic responses without triggering pain. A molecular framework has emerged in which modulating the lifetime of the TrkA:ligand complex can lead to functional selection of downstream signaling pathways. This mechanism of action could spur the development of NGF^painless^-inspired biologics that selectively block nociception in chronic pain^13,16–18^ or enhance the trophic and neuroprotective roles of TrkA:NGF in neurodegenerative conditions^31,49–53^. This general concept of modulating the kinetic parameters of receptor:ligand functional complexes could be extended to bias signaling in other membrane proteins that simultaneously trigger distinct pathways.

## Results

### Activation of TrkA by NGF^painless^ fails to sensitize TRPV1 to capsaicin

Prhevious studies have shown that when TrkA is activated by NGF, it sensitizes TRPV1 channels to stimulation by capsaicin or other noxious stimuli^43–45^. PLCγ1 activation by TrkA and subsequent phosphatidylinositol 4,5-bisphosphate (PIP_2_) hydrolysis have been implicated to play crucial roles in this sensitization^44,46–48^. We, therefore, tested the effectiveness of NGF^painless^ in sensitizing TRPV1 to capsaicin stimuli.

We generated HEK293 stable cell lines co-expressing TrkA-GFP and untagged TRPV1, reported by an mCherry expressed from an independent cassette in the same lentiviral vector (**Fig. 1a**). In a whole-cell configuration, we clamped membrane potential at –60 mV and recorded current responses under different treatment conditions. Capsaicin (10 μM) was first applied as a 1 sec pulse at 30 sec intervals until stable current responses were obtained (**Fig. 1b**, control). Cells were then perfused with 100 ng/mL of either NGF or NGF^painless^ and incubated at 25°C for 15 min. Following incubation, capsaicin-evoked currents were recorded from the same cells using the same capsaicin stimulation protocol as above (**Fig. 1b, Supplementary Fig. 2**).

Consistent with TRPV1 sensitization, NGF treatment significantly increased capsaicin-evoked currents in cells compared to the capsaicin treatments alone (**Fig. 1b-d, Supplementary Fig. 2**). In contrast, NGF^painless^ treatment showed only a minuscule increase in capsaicin-evoked currents (**Fig.1b-d, Supplementary Fig. 2**). This data demonstrates that when TrkA is activated by NGF^painless^, it fails to effectively sensitize TRPV1 channels to capsaicin, disengaging TrkA from the nociceptive signaling pathways.

### NGF^painless^ biases TrkA downstream signaling to disengage nociceptive pathways

When NGF binds to and activates TrkA, it triggers the activation of three downstream proteins, ERK, AKT, and PLCγ1 (**Fig. 2a**). To understand the molecular basis of the selective loss of nociceptive TrkA signaling by NGF^painless^, we assessed whether phosphorylation levels of ERK, AKT, and PLCγ1 change upon stimulation of SHSY5Y cells expressing TrkA with saturating amounts of (50 ng/mL) NGF or NGF^painless^ for 15 minutes.

**Figure 2:**
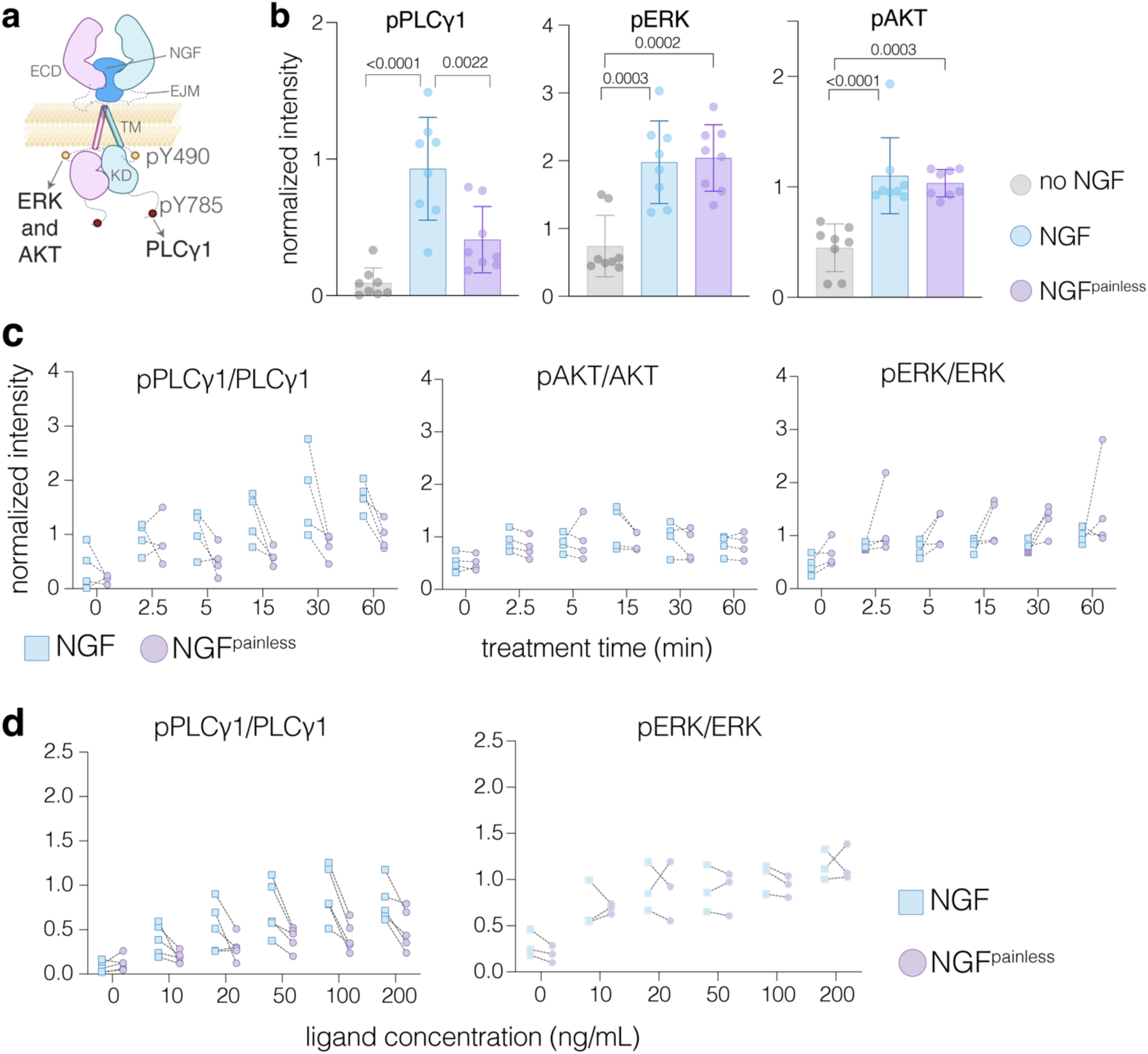
Biased selection of TrkA downstream signaling pathways. **a.** Cartoon depicting the overall domain organization of the heterotetrameric 2:2 complex between the receptor tyrosine kinase, TrkA, and nerve growth factor, NGF. The two autophosphorylation sites on TrkA, Y490 and Y785, that recruit and activate the highlighted downstream signaling pathways, are shown. ECD - extracellular domain; TM - transmembrane; KD - kinase domain. **b.** Quantification of western blots comparing levels of phosphorylation in PLCγ1, ERK, and AKT following stimulation of TrkA-expressing SHSY5Y cells with NGF (light blue) or NGF^painless^ (purple). No NGF treatment (gray) was used as a control. Phosphorylation levels were quantified using band densitometry and normalized to loading controls using Fiji^61^. Data are shown as mean ± s.d. from N = 8 independent experiments. Results of one-way ANOVA analysis and corresponding P-values are reported. The P-values defined as: * , <0.05; **, <0.1; ***, <0.001; ****, <0.0001. **c.** Quantification of time-course western blot data for pPLCγ1, pAKT, and pERK (N=4) is shown by lines connecting paired, simultaneously processed samples under NGF or NGF^painless^ treatment over a range of treatment times (0 - 60 minutes). **d.** Quantification of western blot data for the levels of pPLCγ1 (N=5) and pERK (N=3) over six different NGF or NGF^painless^ concentrations (ranging from 0-200 ng/mL). Data are shown by lines connecting paired, simultaneously processed samples under NGF or NGF^painless^ treatment over a range of ligand concentrations. For (c-d), phosphorylation levels for each protein were quantified using band densitometry, and the phosphorylation signal was normalized with respect to the total signal for the corresponding proteins (each normalized first with respect to loading controls).

Immunoblotting analysis with phospho-specific antibodies against ERK (pT202/pY204), AKT (pS473), and PLCγ1 (pY783) revealed that, compared to NGF, NGF^painless^ treatment showed significantly reduced phosphorylation and activation of PLCγ1 (**Fig. 2b, Supplementary Fig. 3b**). However, no such differences were observed in the phosphorylation levels of AKT or ERK (**Fig. 2b**). Overall levels of TrkA or PLCγ1 remain unchanged (**Supplementary Fig. 3a-b**). We also noted a reduction of TrkA autophosphorylation at the two reported sites (pY490 and pY785) upon treatment with NGF^painless^ (**Supplementary Fig. 3a-b**). Despite reduced pY490, the maintenance of ERK and AKT phosphorylation by NGF^painless^ could be explained by compensatory signal amplification in the signaling cascade upstream of these proteins^88,89^. By contrast, PLCγ1 binds directly to TrkA at pY785 and is phosphorylated by TrkA. Thus, its activation is more vulnerable to reduction in pY785^5,87^.

The phosphorylation levels of PLCγ1, ERK, and AKT as a function of time, under these conditions, recapitulated the single time point immunoblotting data (**Fig. 2c**). pPLCγ1 levels were consistently lower at each time point when treated with NGF^painless^. In contrast, pAKT levels remain unchanged, with some samples showing even more ERK phosphorylation upon NGF^painless^ treatment (**Fig. 2c, Supplementary Fig. 3c**). This indicates that NGF^painless^ biases TrkA signaling to disfavor PLCγ1 activation irrespective of the treatment times. We also determined phosphorylation levels of PLCγ1 and ERK in response to varying concentrations of NGF variants. NGF^painless^ consistently yielded lower pPLCγ1 levels across various doses of ligand, unlike pERK levels that were similar for both variants (**Fig. 2d, Supplementary Fig. 3d**). Taken together, the data show that when NGF^painless^ activates TrkA, it selectively reduces direct phosphorylation and activation of PLCγ1 without affecting ERK or AKT activation (**Fig. 2**). This biased signaling to disfavor PLCγ1 activation is consistent with inefficient sensitization of TRPV1 channels to noxious stimuli, thereby disengaging TrkA signaling from nociceptive pathways^44,46–48^.

### NGF^painless^ leads to reduced intracellular calcium release

NGF stimulation of TrkA activates PLCγ1 and subsequently elevates intracellular calcium levels^54,55^. Activated PLCγ1 hydrolyses phosphatidylinositol 4,5-bisphosphate (PIP_2_), generating inositol triphosphate (IP_3_). IP_3_ binds endoplasmic reticulum-resident IP_3_ receptors, which are ligand-gated calcium channels. This triggers intracellular Ca^2+^ release^54^, a well-known driver of peripheral pain perception^56–59^. To evaluate the downstream consequences of reduced PLCγ1 activation by NGF^painless^, we compared the intracellular Ca^2+^ levels in cells treated with NGF or NGF^painless^. In SHSY5Y cells stably expressing TrkA and GCaMP8, a genetically encoded calcium sensor^60^, we quantified the normalized increase in fluorescence (ΔF/F₀, 488/510 nm, **Methods**) upon treatment with either NGF or NGF^painless^ as a measure of Ca^2+^ levels (**Fig. 3a-b**). We observed that NGF^painless^ treatment failed to elevate cytosolic Ca^2+^ to a similar extent as NGF, even at saturating concentrations. The peak amplitude of the ΔF histogram was reduced by half in cells treated with NGF^painless^ compared to NGF (**Fig. 3b**, **Supplementary Fig. 4**). These results show that reduced PLCγ1 activation by NGF^painless^ dampens the release of Ca²⁺ from intracellular stores.

**Figure 3:**
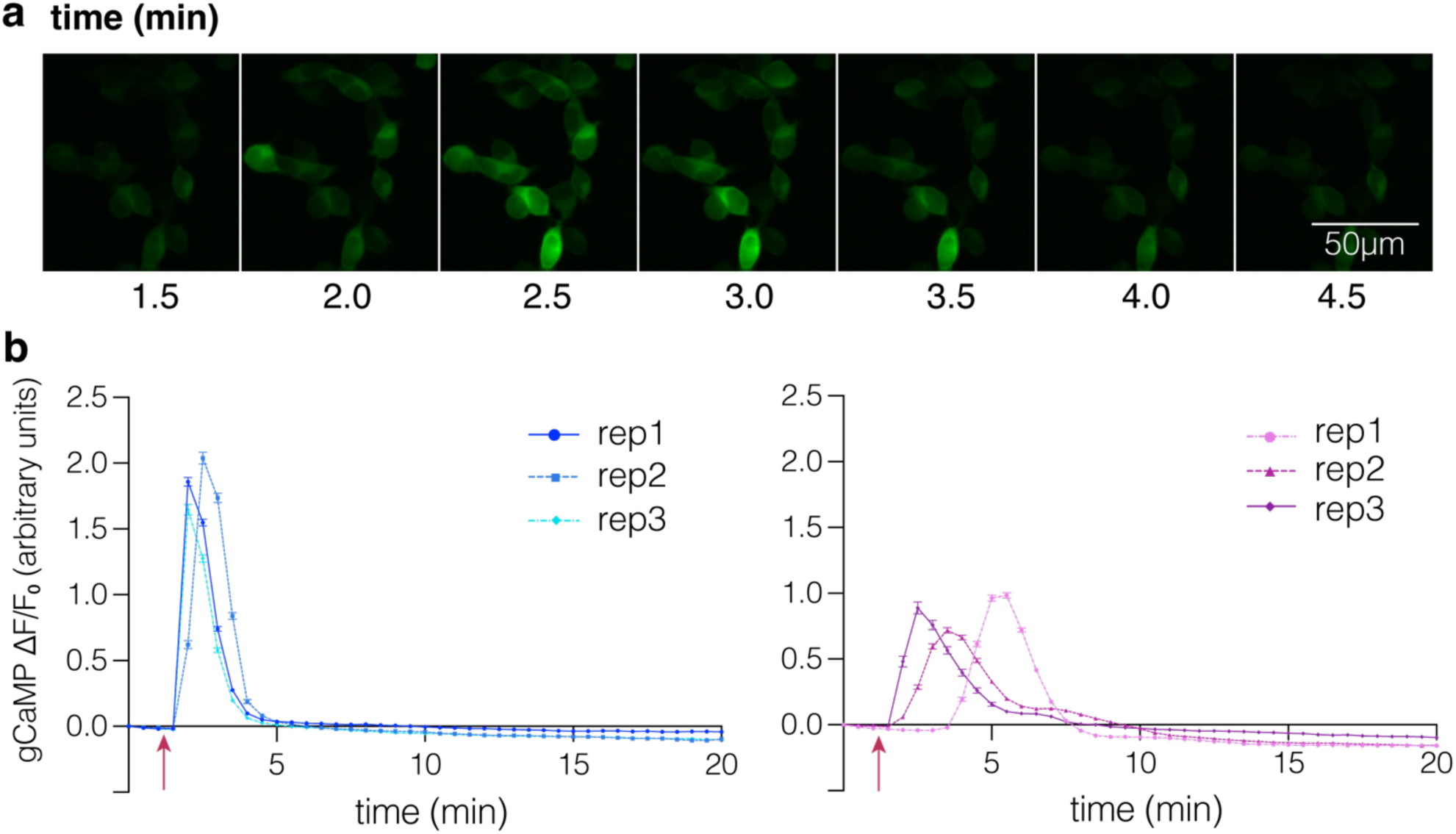
Quantification of intracellular Ca^2+^ levels using GCaMP8 biosensor. **a.** Representative images showing the spike in GCaMP fluorescence upon treatment of TrkA-expressing SHSY5Y cells with a saturating concentration of NGF (100 ng/mL). Scale bar, 50 μm. **b.** Normalized changes in the GCaMP8 fluorescence (ΔF/F₀, 488/510 nm) upon calcium release from intracellular stores in response to 100 ng/mL NGF (top) or NGF^painless^ (bottom) treatment are shown over a period of 20 min. The red arrow indicates the addition of NGF or NGF^painless^. The GCaMP intensity is averaged for 1000 - 2000 single cells (shown as mean ± s.e.m) in each of N = 3 independent experiments.

### NGF^painless^ alters TrkA dynamics on live cell membranes

NGF binding leads to TrkA dimer/multimerization and activation, which in turn phosphorylates and activates downstream effector pathways^62^. To assess whether the native organization of TrkA changes upon binding to NGF or NGF^painless^, we performed single-particle tracking (SPT) analysis of TrkA on live cell plasma membranes^63–66^. In SHSY5Y cells stably expressing near-endogenous levels of ALFA-tagged TrkA-GFP, we labelled TrkA receptors with an anti-ALFA nanobody conjugated to Alexa-647 dye (AF647)^67^. Photobleaching of organic dyes limits the number and length of tracking trajectories, statistically constraining SPT precision and introducing bias in diffusion coefficient estimates^68,69^. To increase the number of trajectories with adequate lengths to compute reliable estimates of diffusion coefficients, we used a ‘photoblueing’ approach of the AF647 dye using a 640 nm laser^70^ (**Fig. 4a, Methods**). The enhanced photostability of photoblued AF647 at 561 nm illumination yielded trajectories extending up to 705 frames, which is equivalent to 38.5 seconds at a frame rate of 20 Hz (**Fig. 4c-d**). We could thereby select ∼100-200 trajectories per cell with more than 100 frames in our single-particle tracking analysis (**Fig. 4b**). Photoblued AF647 also demonstrated spontaneous recovery during prolonged imaging^70^, which allowed us to acquire long TrkA trajectories from the same cell before and after treatment with NGF or NGF^painless^.

**Figure 4:**
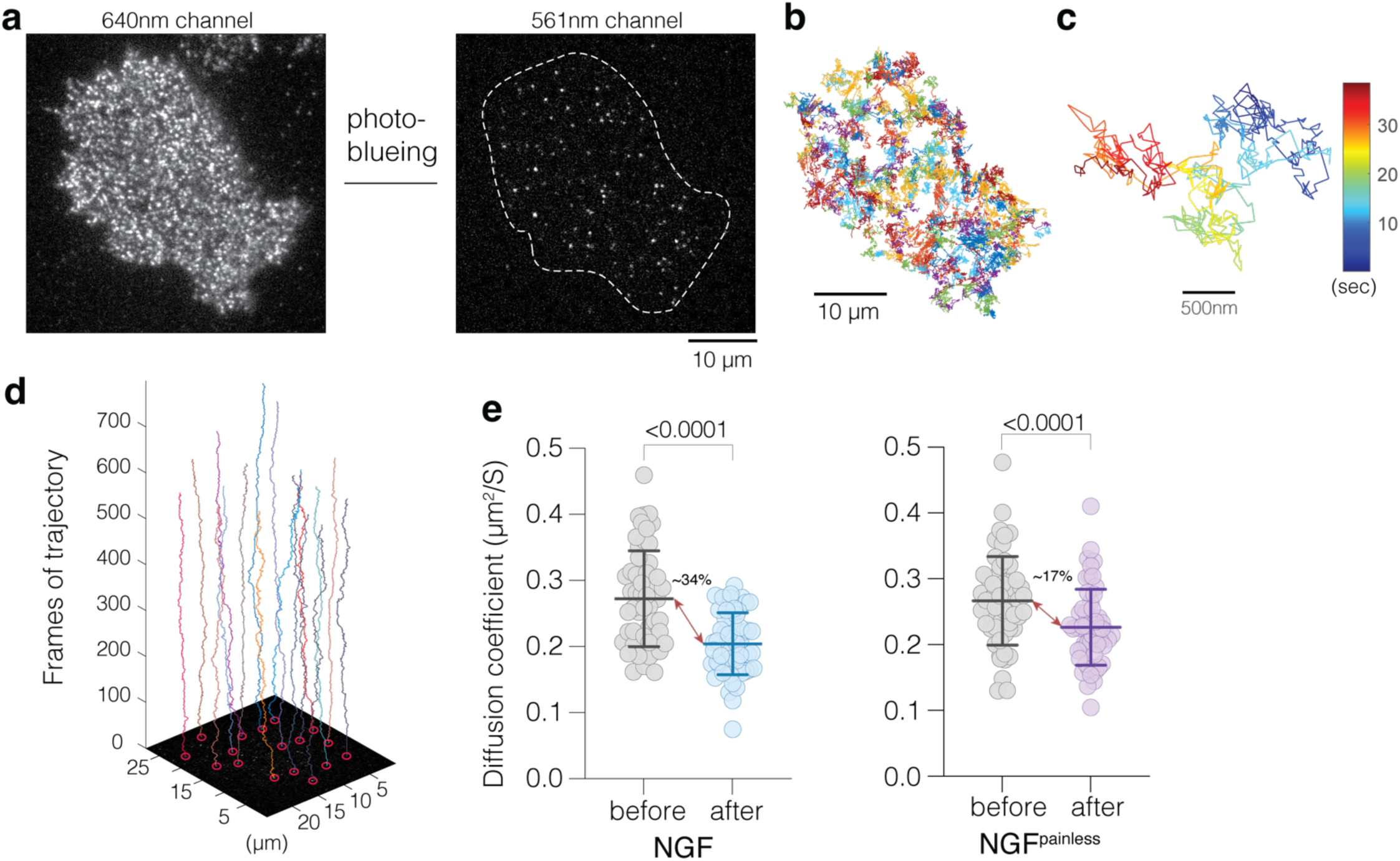
Single-Particle Tracking (SPT) analyses of TrkA lateral dynamics on live SHSY5Y cell plasma membranes. **a.** Cell surface of cells stably expressing ALFA-TrkA-GFP, labeled with anti-ALFA-AF647, photobleached using a 640 nm laser (left panel), producing photoblued AF647-labelled TrkA as visualized with 561 nm laser illumination (right panel; white dashed line indicates cell boundary as detected based on TrkA-GFP signal). Scale bar, 10 µm. **b.** Trajectory map from a representative single SHSY5Y cell featuring a total of 251 long tracks (>100 frames each). Scale bar, 10 µm. **c.** An example of a long trajectory spanning 705 frames (38.5 s at a frame rate of 20 Hz) is shown, with a rainbow color gradient denoting elapsed time in seconds. Scale bar, 500 nm. **d.** An example 3D plot illustrating multiple TrkA trajectories over time (vertical axis, frames) and X–Y axes shows single-particle tracks represented in distinct colors. **e.** Comparison of mean diffusion coefficients for TrkA receptors from single cells before (grey circles) and after (blue circles) treatment with NGF (left) or NGF^painless^ (purple circles, right), respectively. N=56 (left), and N=54 (right). The % difference in the mean diffusion coefficients before and after treatment with NGF or NGF^painless^ is reported. The data is shown as mean ± s.d., and two-sided paired t-test-derived P values are reported.

The reduction in the lateral diffusion coefficient of ligand-bound TrkA is a readout of receptor dimer/multimerization on live cell membranes, as previously reported (**Fig. 4e, Supplementary Fig. 5a**)^63,64,71,72^. We compared the mean diffusion coefficients (κ) of the trajectories formed by tracking single TrkA molecules before and after adding saturating concentrations of NGF or NGF^painless^. NGF significantly reduced κ by ∼34% (N = 56 cells; **Fig. 4e**) while NGF^painless^ reduced κ by only ∼17% (N = 54 cells). This modest but significant difference before and after ligand binding can be explained by the abundance of ligand-independent TrkA dimers on cell membranes^62,73,74^. Unlike other receptor tyrosine kinases, such as EGFR, which form dimers/multimers when a ligand binds^64,72,75^, the changes in κ here only reflect the ligand-induced changes to a pre-existing distribution of TrkA monomers and dimers.

To assess how ligand binding affects the lateral dynamics of TrkA, or the area it explores on cell membranes, we quantified the average length of the major axis of trajectories (**Supplementary Fig. 5b**). NGF binding reduced the average major axis length, suggesting higher spatial constraints on TrkA diffusion, while NGF^painless^ binding induced no discernible change. This data also indicates that NGF reduces the lateral diffusion of ligand-bound TrkA on the plasma membrane more effectively than NGF^painless^, likely because the complex forms a higher proportion of dimers or multimers. This inability of NGF^painless^ to enhance TrkA dimer/multimers effectively is consistent with the ineffective activation of PLCγ1 and its subsequent downstream signaling.

In addition to receptor dimer/multimerization, the reduced diffusion of ligand-bound TrkA can potentially arise from other factors, such as TrkA interaction with lipid microdomains, recruitment of cytoplasmic factors, or interactions with cytoskeletal components^71,76^. Therefore, using Native-nanoBleach (NNB), we directly determined the oligomeric distribution of TrkA upon binding to either NGF or NGF^painless62^. NNB is a single-molecule imaging method that quantifies equilibrium oligomeric distribution of membrane proteins under various conditions, in the context of their native membrane environment at 10 nm lateral spatial resolution^62^.

We used amphipathic co-polymers (SMA) to excise circular patches of cell native membranes, denoted native nanodiscs (NND), containing membrane proteins, which can be affinity-purified, as previously described^62^. We isolated and purified GFP-tagged TrkA:NGF complexes from HEK293 cells in ∼10 nm NND. We reasoned that a long-lived TrkA:ligand complex will have a higher likelihood of being captured within a 10 nm disc compared to relatively short-lived ones, which will likely dissociate before excision into an NND. Fluorescence size exclusion chromatography (FSEC) of NNDs containing TrkA-GFP isolated from untreated cells showed two peaks - a prominent right peak with a small left shoulder (**Fig. 5a**). Upon NGF treatment of cells, the size of the left shoulder in the FSEC profile increases, resolving into a distinct peak, indicating an increase of dimers/multimers. However, the amplitude of this left peak upon NGF^painless^ binding is consistently lower compared to NGF, suggesting reduced amounts of dimer/oligomers (**Fig. 5a**).

**Figure 5:**
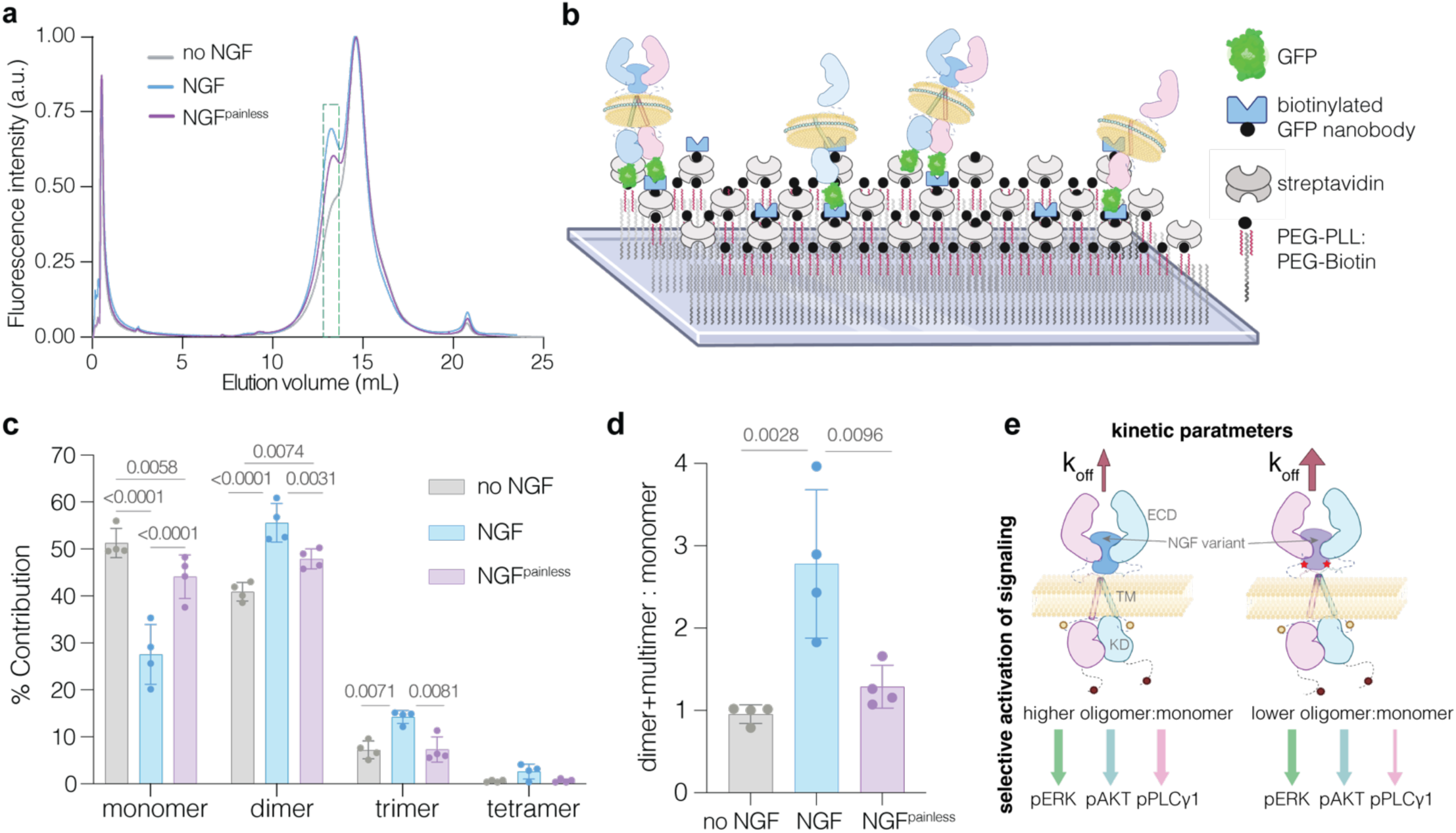
Native-nanoBleach analysis of the equilibrium oligomeric distribution of TrkA:NGF or TrkA:NGF^painless^ signaling complex. **a.** Fluorescence size exclusion chromatography (FSEC) profile of native nanodisc-encapsulated TrkA-GFP isolated from cells treated with either NGF, NGF^painless^, or untreated cells, monitoring C-terminal GFP fluorescence. Treatment with NGF or NGF^painless^ leads to an increase in the amplitude of the left shoulder peak at ∼13.5 ml (highlighted by the green dotted box), indicating an increased population of ligand-induced oligomers. FSEC traces are shown after standard min-max normalization, with the right peak set to 1, for ease of comparison. **b.** Depiction of the setup for Native-nanoBleach, a TIRF-based single-molecule GFP-photobleaching step analysis. The cartoon depicts the functionalization of glass slides (substrates) to display a GFP nanobody bait for capturing GFP-tagged TrkA complexes isolated in native nanodiscs. **c.** Native-nanoBleach analysis of the left peak fractions in (a) from untreated, NGF- or NGF^painless^-stimulated samples shows TrkA oligomeric distribution. Consolidated data from 3,000 - 4,000 single nanodiscs from N = 4 biologically independent samples represented as mean ± s.d. Results of two-way ANOVA analysis with P-values are reported. **d.** Dimer+multimer (oligomer) to monomer ratio in native nanodiscs for the untreated or NGF/NGF^painless^-bound TrkA calculated from (c). N = 4 biologically independent samples represented as mean ± s.d. Results of two-way ANOVA analysis with P-values are reported. For (c-d), **** for P < 0.0001, *** for P <= 0.001, ** for P <= 0.01, * for P <= 0.05 **e.** Schematic showing the relationships among TrkA:NGF/NGF^painless^ complex lifetime, which is inversely proportional to k_off_, the ratio of oligomer to monomer on native membranes, and how this may influence selective downstream signaling.

To unambiguously determine the equilibrium oligomeric distribution of the different ligand-bound states of TrkA, we immobilized the fractions, corresponding to the non-void, left peak of FSEC, on glass slides using GFP-nanobody baits (**Fig. 5b, Supplementary Fig. 5c**). We then counted the number of TrkA subunits within single NNDs isolated from mock-treated cells or those treated with saturating amounts of NGF or NGF^painless^ (**Supplementary Fig. 5d-e**)^62^. Unliganded TrkA showed ∼1:1 dimers+multimers (oligomer) to monomer ratio (**Fig. 5c-d**)^62^. NGF binding increases this ratio by ∼3-fold (**Fig. 5c-d**), while NGF^painless^ showed only a moderate increase by ∼1.4-fold. So, the TrkA:NGF^painless^ complex has a ∼2-fold lower abundance of signaling-competent receptor dimers/multimers at equilibrium, even at saturating concentrations of ligand (**Fig. 5c-d)**. These data demonstrate that the TrkA:NGF^painless^ complex has a lower lifetime compared to that of TrkA:NGF on native membranes, leading to functional selection of the ERK and AKT pathways and biased reduction of PLCγ1 pathway activation (**Fig. 5e**).

### Perturbed electrostatic complementarity at the TrkA:ligand interface lowers TrkA:NGF^painless^ lifetime

To determine whether NGF^painless^ binding alters the overall architecture of 2:2 TrkA:ligand complex, we solved the structures of the full-length TrkA:NGF or TrkA:NGF^painless^ complexes by single particle cryo-EM (**Fig. 6a-d, Supplementary Fig. 6-9, Supplementary Table 1**). For these structural studies, we purified TrkA:NGF or TrkA:NGF^painless^ complexes from Expi293 cells and reconstituted them in lipid nanodiscs that mimic the plasma membrane environment (**Supplementary Fig. 6a, b and Supplementary Fig. 7a, b)**. As noted previously with cryo-EM studies of other single-pass transmembrane receptor tyrosine kinases^77–80^, high-resolution reconstruction with simultaneously well-resolved extracellular domains (ECD) and transmembrane (TM) / kinase domains (KD) was not feasible (**Supplementary Fig. 10a-b**). Nevertheless, we were able to resolve the 2:2 complexes of NGF or NGF^painless^ bound to the TrkA ECDs, specifically of the NGF and TrkA interaction interface (ECD-Ig2) at 3.79 Å and 2.87 Å, respectively (**Fig. 6a-b, Supplementary Fig. 8, 9**). This resolution allowed us to compare and understand the differences in structures and dynamics between the two TrkA:ligand interfaces within the intact signaling complex.

**Figure 6:**
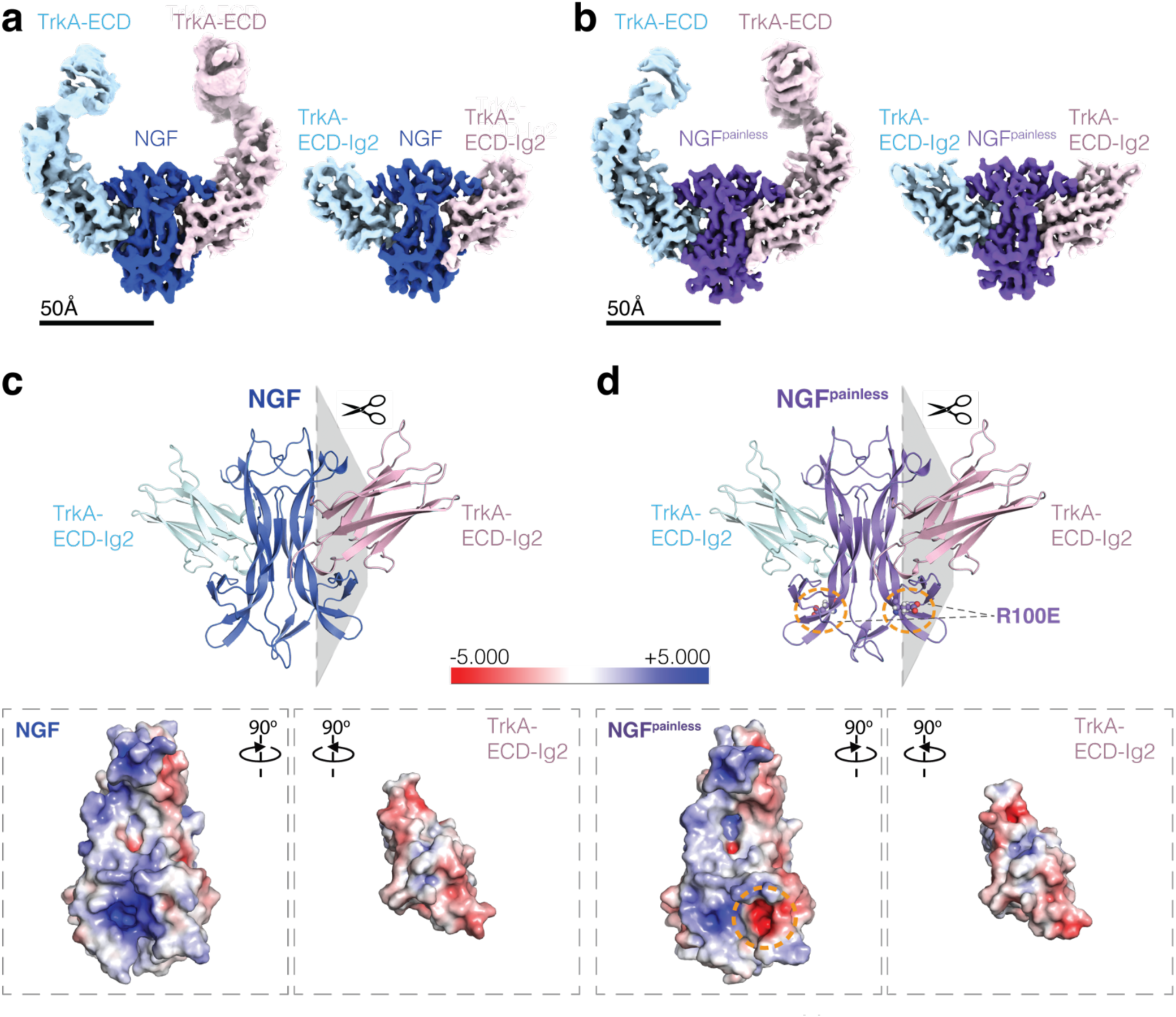
Structural analysis of the 2:2 TrkA:NGF and TrkA:NGF^painless^ complexes. **a.** Top panel, left: 4.02 Å Coulomb potential map of the 2:2 TrkA:NGF complex, with individual TrkA-ECD monomer densities shown in cyan and pink, and the dimeric-NGF shown in blue. Right: Inset shows Coulomb potential map of the TrkA-ECD-Ig2:NGF core region forming the receptor-ligand interface at 3.79 Å. Scale bar, 50 Å. **b.** Top panel, left: 3.13 Å Coulomb potential map of the 2:2 TrkA:NGF^painless^ complex, with individual TrkA-ECD monomer densities shown in cyan and pink, and the dimeric-NGF^painless^ shown in purple. Right: Inset shows Coulomb potential map of the TrkA-ECD-Ig2:NGF^painless^ core region forming the receptor-ligand interface at 2.87 Å. Scale bar, 50 Å. **c.** Top: deposited model of the TrkA-ECD-Ig2:NGF core (PDB ID: YKU). Bottom: Surface electrostatic potential rendering of the NGF and TrkA ECD-Ig2 domains, showing the interaction interface. An adaptive Poisson-Boltzmann Solver was used to calculate this electrostatic potential after isolating the NGF and the contacting TrkA ECD-Ig2 arm, as depicted by the gray plane. The interface shows electrostatic complementarity that results in the formation of the TrkA:NGF complex. **d.** Top: deposited model of the TrkA-ECD-Ig2:NGF^painless^ core (PDB ID: YKT). Bottom: Surface electrostatic potential rendering of the NGF^painless^ and TrkA ECD-Ig2 domains, showing the interaction interface. An adaptive Poisson-Boltzmann Solver was used to calculate this electrostatic potential after isolating the NGF^painless^ and the contacting TrkA ECD-Ig2 arm, as depicted by the gray plane. The electrostatic complementarity is perturbed due to the lowering of the net positive charge on the NGF^painless^ interface from the charge reversal (R100E) mutation. The site of mutation is labeled, highlighted as spheres, and encircled by orange dotted lines on the cartoon and Coulomb potential map. The APBS colormap is scaled equally for (c-d).

We observed that the overall architecture of TrkA:NGF and TrkA:NGF^painless^ dimers was nearly identical (**Fig. 6a–b**). To probe potential differences, we examined the TrkA:ligand interface by calculating the surface electrostatic potential of NGF and the TrkA ECD-Ig2 domain in the 2:2 complex using Adaptive Poisson–Boltzmann methods^81^. The NGF binding surface exhibited a broad electropositive patch that is complementary to the electronegative surface of TrkA ECD-Ig2 (**Fig. 6c**). This charge complementarity provides multiple opportunities for TrkA ECD-Ig2 to engage a large accessible surface on NGF, thereby supporting formation of a stable and dynamic 2:2 complex. In NGF^painless^, the charge reversal R100E mutation reduces the net electropositive potential of the TrkA-binding surface, diminishing the electrostatic complementarity at the interface (**Fig. 6d**). As a result, it may spatially restrict how NGF^painless^ engages the negatively charged ECD-Ig2 arm, limiting the range of accessible conformations. We propose that this constraint introduces a strain within the TrkA:NGF^painless^ complex, increasing its off-rate, which in turn reduces its lifetime on membranes. Importantly, in all our reconstructions, we did not impose a C2-symmetry, so that the dynamic conformational asymmetry is retained for the individual ECD arms.

In both complexes, the local resolution decreases progressively from the core, comprising the NGF and the TrkA-Ig2 domains, toward the extended ECD arms (**Supplementary Fig. 10c-d**). This gradient is more pronounced in the ECD arms of the TrkA:NGF complex than in TrkA:NGF^painless^, where both arms are better resolved, with one slightly more so than the other. The better resolution of the ECD arms in TrkA:NGF^painless^ is consistent with higher spatial constraints from reduced electrostatic complementarity, which may limit the conformational sampling of the ECD at the TrkA:NGF^painless^ interface compared to TrkA:NGF. To exclude particle-number bias, we chose two random particle subsets, each with about half the number of total particles in the TrkA:NGF^painless^ dataset, repeated the analysis, obtaining comparable results each time (**Supplementary Fig. 10e**).

### Role of the disordered extracellular juxtamembrane regions in regulating TrkA:ligand complex lifetime

We noted that the sequence of the extracellular juxtamembrane (EJM) region of TrkA immediately following the last resolved residue in our model is largely negatively charged, with several conserved glutamic acid (E384, E388) and aspartic acid (D389) residues (numbering based on Uniprot ID: P04629). We used Alphafold 3 (AF3) to evaluate the positioning of the first 15 EJM residues with respect to the TrkA-ECD:NGF interface (ipTM = 0.68; pTM = 0.7; **Fig. 7a**). The predicted model aligned well with our high-resolution structure of TrkA-ECD-Ig2:NGF (RMSD of ∼0.9 Å). We noted that the electrostatic complementarity observed at the TrkA:NGF interface continued with this EJM region, with negatively charged EJM residues cradling the lower part (β7-β8) of NGF (**Fig. 7a**). NGF^painless^ lowered the net electropositive charge in this region (**Fig. 6d**), possibly disrupting these predicted interactions with the EJM.

**Figure 7:**
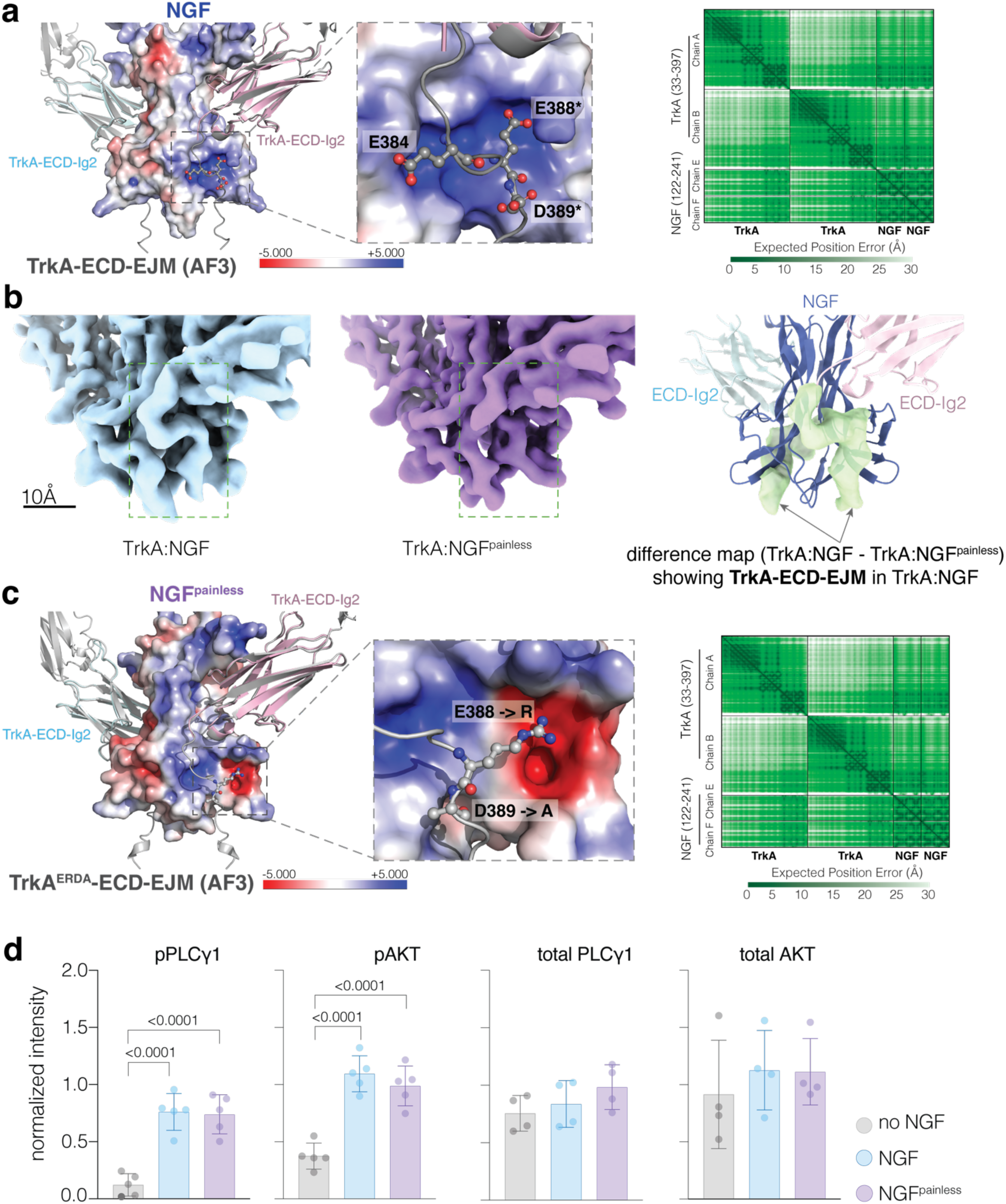
Role of TrkA extracellular juxtamembrane region (EJM) in NGF engagement and downstream signaling. **a.** Left. Alphafold3 prediction (gray) of the TrkA:NGF 2:2 complex, highlighting the Ig2 of TrkA-ECD and the first 15 residues of the EJM. The surface electrostatic potential is shown for the AF3-predicted NGF dimer. The corresponding experimentally obtained high-resolution TrkA-ECD-Ig2 subunits are aligned on the AF3 model (gray) and are colored in pink and cyan, respectively. A zoomed-in inset shows the specific EJM residues participating in the electrostatic interaction with β7-β8 of NGF (the numbering of TrkA and NGF based on UNIPROT IDs: P04629 and P01138, respectively). Right. Heatmap showing expected position error (in Å) in the prediction. **b.** Left: Close-up view of the extracellular juxtamembrane (EJM) region of TrkA:NGF complex, EMD-73058. Middle: Close-up view of the extracellular juxtamembrane (EJM) region of TrkA:NGF^painless^ complex, EMD-73057. All maps are low-pass filtered to 5 Å, resampled, and presented at the same threshold. Right: Additional density present in the TrkA:NGF map for ∼10 EJM residues, which is absent in TrkA:NGF^painless^ map, is shown in green. The deposited model, featuring ECD-Ig2 and NGF, is shown with a similar color scheme to that in Supplementary Figure 8, for reference. Scale bar, 10 Å. **c.** Left. Alphafold3 prediction (gray) of the TrkA^ERDA^:NGF^painless^ 2:2 complex, comprising the ECD and the mutated EJM. The corresponding experimentally obtained high-resolution TrkA-ECD-Ig2 subunits from the TrkA:NGF^painless^ complex are aligned on the AF3 model (gray) and are colored in pink and cyan, respectively. The surface electrostatic potential is shown for the experimentally obtained NGF^painless^ dimer. Zoomed-in inset highlights the specific EJM residues that were mutated in TrkA^ERDA^ (numbering of TrkA and NGF based on UNIPROT IDs: P04629 and P01138, respectively; mutations also highlighted by * in the zoomed-in inset of panel a). The APBS colormap is the same as in (a). Right. Heatmap showing expected position error (in Å) in this prediction. **d.** Quantification of western blots comparing the levels of phosphorylation in PLCγ1 and AKT, along with total levels of PLCγ1 and AKT, following stimulation of TrkA^ERDA^ with NGF (light blue) or NGF^painless^ (purple). No NGF treatment (gray) was used as a control.

This observation is strengthened by the presence of low-resolution density for the first ∼10 EJM residues in the TrkA:NGF complex map, which is absent from TrkA:NGF^painless^ (**Fig. 7b**). This additional density supports the AF3-proposed interactions between EJM and NGF (**Fig. 7a**), which allows the flexible EJM region to be partially resolved in the TrkA:NGF complex. While past studies have proposed the importance of the EJM in NGF binding and the regulation of TrkA signaling^82,83^, our data provide the first structural evidence of such an interaction between EJM and NGF. In the TrkA:NGF^painless^ complex, the repulsion between the negatively charged EJM and the negatively charged pocket at the base of NGF^painless^, introduced by the R100E charge reversal mutation (**Fig. 6d**), likely disrupts the EJM interactions. This disruption could explain the absence of EJM density in the map (**Fig. 7a, b**). These results reveal a structural mechanism in which the electrostatic complementarity between the structured ECD-Ig2, along with the unstructured EJM of TrkA and the ligand, together dictates the TrkA:NGF complex lifetime, which, in turn, modulates downstream signaling. This complementarity is disrupted with NGF^painless^, explaining the lowered lifetime of the TrkA:NGF^painless^ complex compared to TrkA:NGF and the resulting biased signaling.

### Structure-guided charge reversal mutations in TrkA-EJM rescues PLC**γ**1 phosphorylation by NGF^painless^

Our data suggest that the perturbed electrostatic complementarity between TrkA-ECD-Ig2 plus EJM and NGF^painless^ lowers complex lifetime, leading to a biased reduction in PLCγ1 phosphorylation. We further validated this by introducing two point mutations in TrkA-EJM (E388R and D389A; hereafter called TrkA^ERDA^), which collectively reduce the overall negative charge of the EJM region and restore the electrostatic complementarity between EJM and NGF^painless^ (**Fig. 7a, c, zoomed inset**). Moreover, an AF3 model of the 2:2 TrkA^ERDA^:NGF^painless^ complex positioned the mutated E388 -> R residue to form a hydrogen bond with E100, compensating for the R100E charge reversal mutation in NGF^painless^ (ipTM = 0.68; pTM = 0.69; **Fig. 7c**). We propose that TrkA^ERDA^ will restore the PLCγ1 phosphorylation by NGF^painless^ to the similar levels as TrkA:NGF.

To test this, we treated SHSY5Y cells expressing the mutant TrkA^ERDA^ with NGF^painless^ or NGF for 15 min and quantified the phosphorylation levels of PLCγ1 along with AKT as control. TrkA^ERDA^ shows comparable phosphorylation levels for both PLCγ1 and AKT when activated by either NGF or NGF^painless^ (**Fig. 7d, Supplementary Fig. 3e**). The total levels of PLCγ1 and AKT remain unchanged between the treatment conditions (**Fig. 7d**). This rescued PLCγ1 phosphorylation in TrkA^ERDA^ upon treatment with NGF^painless^ firmly established the importance of the EJM region in expanding the TrkA:NGF/NGF^painless^ interface beyond the structured domains alone. This data provides the first structural evidence of the TrkA-EJM region interaction with NGF in addition to ECD-Ig2. Our results underscore the importance of modulating these interactions, as observed with NGF^painless^, to adjust the TrkA:ligand complex lifetime and bias downstream signaling.

## Discussion

The TrkA:NGF pathway is a highly sought-after therapeutic target. It has been targeted with agonists to promote neurogenesis and with antagonists, such as anti-NGF antibodies, to relieve pain^13,17,50,51,84,85^. However, complete stimulation or inhibition of TrkA has been limited clinically by several adverse side effects. The agonists promote neuronal growth, but induce pain and hyperalgesia, while the antagonists alleviate pain but cause neurological issues and accelerate joint damage^10,12–14,17,18,51,86^. Hence, there is an urgent need to understand how to selectively modulate TrkA downstream signaling, rather than simply activating or inhibiting it overall. Using NGF^painless^, an NGF variant inspired by rare mutations in humans that are impervious to pain, we showed how to uncouple the neurotrophic and nociceptive TrkA signaling. This mechanistic knowledge provides a framework to engineer next-generation biologics that selectively relieve pain while minimizing neuronal adverse effects.

TrkA oligomerization is critical for its autophosphorylation and downstream signaling^5,6,87^. Changing the lifetime of the TrkA:ligand signaling complex can influence the extent of TrkA autophosphorylation and lead to selective activation of downstream pathways (**Fig. 5e**). We show that for NGF^painless^, the lifetime of the TrkA:ligand complex is shortened, such that it reduces autophosphorylation at both Y490 and Y785 on TrkA (**Supplementary Fig. 3a-b**). However, NGF^painless^ biases signaling downstream of TrkA – ERK and AKT activation are maintained, whereas PLCγ1 signaling and subsequent calcium release are attenuated (**Figs. 2, 3**). This could be explained by the fact that ERK and AKT activation result from several sequential signaling steps downstream of phospho-Y490^5,87^, which provide multiple opportunities to buffer a small reduction in autophosphorylation^88,89^. By contrast, PLCγ1 directly binds to TrkA at pY785 and is phosphorylated by TrkA to facilitate its activation, with no intermediate steps. Thus, its activation is likely more vulnerable to reduced TrkA autophosphorylation^5,87^.

We show that, unlike NGF, NGF^painless^ fails to sensitize TRPV1 channels (**Fig. 1**), explaining the reduced nociception observed in the genetic condition, HSANV, and in animal models of NGF^painless^. Previous studies have reported that the activation of PLCγ1 and PI3K via TrkA sensitizes TRPV1^43,44^. We unequivocally demonstrate that NGF^painless^ reduces phosphorylation and activation of only PLCγ1 without altering ERK or AKT signaling. Our results suggest that following the activation of TrkA, the extent of PLCγ1 activation and signaling is a critical determinant of TRPV1 sensitization.

The structures of TrkA in complex with NGF or NGF^painless^ reveal that electrostatic complementarity at the TrkA:ligand interface, which extends into the TrkA-EJM, regulates the lifetime of the complex in native cell membranes (**Figs. 6, 7**). Our cryo-EM map of TrkA:NGF provides first structural evidence, albeit low-resolution, for the first ∼10 residues in the EJM linker, which cradles the electropositive surface at the lower part of NGF. An AF3 model prediction supports this, highlighting the EJM interacting with NGF and contributing to its engagement by TrkA, which may likely influence TM domain positioning and the coupling of the ECD with the KD. Our data provide a mechanistic basis for previous studies that show that mutations or truncations in TrkA-EJM result in attenuated NGF-mediated response^82,83^. In contrast, NGF^painless^ perturbs this overall electrostatic complementarity and likely repels the negatively-charged EJM, leading to shorter TrkA:NGF^painless^ complex lifetime on cell membranes (**Figs. 4, 5**). This is further attested by the lack of EJM density in the TrkA:NGF^painless^ map despite its overall higher resolution compared to TrkA:NGF (**Fig. 7b**). Finally, we further validated our model by showing that the selective reduction in PLCγ1 activation by NGF^painless^ can be rescued by introducing structure-guided, compensatory mutations in TrkA-EJM that reduce its net negative charge (**Fig. 7c-d**).

Together, our findings reveal that differential electrostatic complementarity at the TrkA:ligand interface governs complex lifetime and biases downstream signaling, providing a molecular basis for uncoupling the neurotrophic and nociceptive outcomes of TrkA activation. Leveraging NGF^painless^, we exposed a tractable vulnerability in the TrkA:NGF complex, opening up new opportunities to engineer biologics that uncouple nociceptive and neurotrophic pathways, allowing for the selective modulation of one without impacting the other. Our work provides a new perspective on an attractive therapeutic target to realize next-generation, non-opioid therapies for chronic pain management.

## Author Contributions

M.B. and S.K. designed the experiments with contributions from H.A.J., G.W., R.F., Y.K., and S.M. S.K., G.W., and M.T.H. performed and analyzed all immunoblotting and calcium signaling data with inputs from M.B. S.K., G.W., and M.T.H. performed single-molecule experiments with support from L.K. S.K., G.W., and M.T.H. analyzed the single-molecule data. S.K. and H.A.J. purified protein and prepared samples for cryo-EM data with inputs from J.K.R. and V.K. H.A.J., J.K.R., V.K., S.M., and M.B. collected, analyzed, and interpreted the cryo-EM and structural data. F.M., S.C., and A.C. provided the NGF^painless^ and helpful experimental suggestions. S.K., J.K., S.J.M., and B.P.H. contributed to the generation and maintenance of various stable cell lines. W.J. performed and analyzed native mass spectrometry experiments. R.F., S.K., and Y.K. prepared samples, performed, and analyzed the electrophysiology experiments. M.B., S.K., S.M., and Y.K. wrote the manuscript with critical contributions from H.A.J., G.W., M.T.H., and co-authors.

## Acknowledgments

We thank members of the Bhattacharyya lab for helpful discussions. We especially thank Rhys Girouard for maintenance of the Bhattacharyya lab microscopy setup and Drs. David and Clotilde Calderwood for access and help with the microscopy setup for the calcium signaling experiments. We thank Dr. Kallol Gupta for helpful suggestions, discussions, and access to the native mass spectrometry instrument in his lab. We thank Dr. Marc Llaguno for help with the maintenance of the cryo-EM facility at the Yale School of Medicine. We thank the staff of the University of Michigan cryo-EM facility for their assistance and help in data collection and microscope maintenance. We thank Dr. Shalini Low-Nam, Purdue University, and Dr. Leonard Kaczmarek, Yale University, for helpful early discussions on single-particle tracking and electrophysiology experiments, respectively. We thank Dr. Neel H. Shah, Columbia University, and Drs. Mark Lemmon and Ben Turk, Yale University, for insightful discussions on receptor signaling. We also thank members of the Department of Pharmacology at Yale University for helpful discussions on this work. Expi293 cells were a gift from Dr. Karin Reinisch’s lab at Yale University, HEK293 cells were a gift from Dr. John Kuriyan’s lab at Vanderbilt University, and SF9 cells were a gift from Dr. Joel Butterwick’s lab at Yale University. NGF was a gift from Genentech, USA. pHRSIN and pEXQV constructs were a gift from Dr. Shalini Low-Nam, Purdue University. All the work reported here was performed in the United States.

## Data availability

Single-particle cryo-EM maps of TrkA:NGF and TrkA:NGF^painless^ have been deposited at the Electron Microscopy Data Bank (EMDB) with the accession codes EMD-73058 and EMD-73057, respectively. The corresponding models have been deposited in the PDB with accession codes 9YKU and 9YKT, respectively. All other materials used and data reported in the study are available upon request.

## Materials and Methods Preparation of plasmids

### Mammalian expression constructs

A list of all constructs generated in this study, including their backbone plasmids and the positions of various tags, is provided in **Supplementary Table 2**. Monomeric enhanced GFP (A206K mutation; referred to as GFP throughout) was used throughout in this study. TrkA and its various tagged versions were cloned into the pEGBacMam-C-term-3C-GFP-His8 plasmid (Addgene, #160687). To generate pEGBacMam-C-term-3C-GFP-3×-FLAG, the His8 tag at the C-terminus was replaced with a 3× FLAG tag using Gibson assembly (New England Biolabs, #E2611S). An Alfa-tag sequence was introduced at the N-terminus of TrkA immediately downstream of the signal peptide in some constructs. The Alfa-tag fragment was PCR-amplified with gene-specific primers and inserted via Gibson assembly. All constructs were verified by Sanger sequencing and the final positive version of the construct used in each study was subjected to Plasmid-EZ Whole Plasmid Sequencing to confirm the sequence integrity of the full construct (Genewiz, Azenta).

### Lentiviral constructs

The TrkA gene and its variants (**Supplementary Table 2**) were cloned into the second-generation lentiviral transfer vector pSA75-pHRSin using Gibson assembly (New England Biolabs, #E2611S). Vector backbone and insert fragments were generated by PCR amplification using Phusion high-fidelity DNA polymerase (New England Biolabs, #M0530S). The amplified PCR products were gel-purified and treated with DpnI to remove the parental plasmid. Gibson assembly reactions were performed using a 1:3 ratio of vector backbone to insert DNA in a 10-15 μL reaction volume and incubated at 50°C for 30 minutes. Assembly products were transformed into NEB® Stable Competent *E. coli* (High Efficiency, NRB, #C3040) and plated on LB-agar selection plates supplemented with ampicillin (100 µg/mL). Plates were incubated at 30°C for 36-48 hours. Individual colonies were screened by Sanger sequencing of junction regions, and a final positive clone was subjected to complete plasmid sequencing to confirm the sequence integrity of the construct (Genewiz, Azenta). The 3^rd^ generation pLV[Exp]-mCherry/Puro-EF1A>hTRPV1[NM_080706.3 (vector ID: VB900181-7765gub) and pLV[Exp]-Puro-EF1A>jGCaMP8m (vector ID: VB900169-8014wfn) was purchased from VectorBuilder. These 2^nd^ and the 3^rd^ generation lentiviral plasmids were used with standard 2^nd^ and 3^rd^ generation helper plasmids, respectively, for the generation of lentivirus as described below.

## Tissue Culture

### Adherent cells

HEK293FT cells were cultured and maintained in DMEM (Gibco, #11965-092) media supplemented with 10% FBS (Sigma-Aldrich, #12306C) and 1% antibiotic-antimycotic (AA, Thermo Fisher Scientific, #15240062). Advanced DMEM/F12 medium (Thermo Fisher Scientific, #12634010) supplemented with 10% FBS, 1mM L-Glutamine (Thermo Fisher Scientific, #25030081), and 1mM AA was used to grow and maintain SHSY5Y cells. For regular maintenance of HEK293FT/SHSY5Y cells, cells were detached using 0.05% trypsin (Thermo Fisher Scientific, #25300-054) at 37 °C for 3-4 minutes, resuspended in media, and plated at a dilution of 1:20. The plates were maintained in an incubator at 37°C with 5% CO2 and 80% humidity. For experiments, cells were split at 1:2 dilution and used the next day.

An SHSY5Y neuroblastoma cell line stably expressing TrkA (Kerafast, #ECP006) was grown and maintained according to the supplier’s instructions. Briefly, cells were detached from the plate surface using 0.05% Trypsin for 3-4 minutes at 37°C and resuspended in RPMI1640 media supplemented with 10% FBS, 1% pen-strep (Thermo Fisher Scientific, #15140122), and 0.3mg/ml G418 (Geneticin, Thermo Fisher Scientific, #10131035). Cell clumps were carefully homogenized via resuspension with a serological pipette. Cells were regularly maintained at 1:10 dilutions and. For experiments, 1:2 dilutions were made, and cells were used the following day.

### Suspension cells

Expi293 cells (a gift from Dr. Karin Reinisch’s lab, Yale University) were maintained in Expi293 expression medium (Thermo Fisher Scientific, #A1435101) at 37°C with 8% CO₂ in an incubator with 80% humidity under constant orbital shaking (120–150 rpm). Cultures were seeded at 0.5 × 10^6^ cells/mL and regularly passaged at a maximum density of 5 million cells/mL to maintain growth. Cell density and viability were routinely monitored using a trypan blue dye. Sf9 cells (ATCC, #CRL-1711) were cultured in Sf-900 II SFM media (Thermo Fisher Scientific, #10902088) at 27°C in a shaking incubator. Cells were mixed with trypan blue dye (Gibco, #15250-061) at a 1:4 ratio (cells:dye) and counted using a hemocytometer. Cells were pelleted at 1000 g for 4 min, supernatant discarded, and the cell pellet was resuspended in fresh medium. Subcultured cells were maintained at 27°C under the same conditions. The media and its components are also summarized in **Supplementary Table 7.**

## Plasmid transfection and transduction

### Transient transfection

We followed the standard polyethylenimine (PEI) protocol for transient transfection of constructs in mammalian cells. For adherent HEK293FT cell lines, plasmid DNA and PEI (3 μl PEI per 1 μg DNA) were separately diluted in Opti-MEM Reduced Serum Medium (Gibco, # 31985070), combined after 5 min, and subsequently incubated for 20 min at room temperature. The transfection mix was subsequently added dropwise to adherent cultures at approximately 80–90% confluency.

### Bacmid preparation and transduction

Bacmid DNA was prepared using *E. coli* DH10Bac cells (Thermo Fisher Scientific, #10361012, **Supplementary Table 4**) and isolated following a previously described protocol^90^. Briefly, 50 ng of plasmid was transformed into DH10Bac competent cells and plated on LB agar selection plates containing kanamycin (50 μg/ml, GoldBio,#K-120-50), gentamicin (7 μg/ml, Gibco, #15710-064), tetracycline (10 μg/ml, Sigma-Aldrich, #T7660-5G), IPTG (0.17 mM, GoldBio, #12481C25), and Bluo-gal (100 μg/ml, RPI, #B72500-1.0). Plates were incubated at 37°C for ∼48 hours in the dark. White colonies were re-streaked on fresh selection plates for confirmation, and these white colonies were used for bacmid DNA extraction. For transfection, Sf9 cells were seeded at 2 × 10⁶ cells/ml in 6-well plates with Sf-900 II SFM media (Thermo Fisher Scientific, #10902088) and incubated at 27°C for 20 min to allow cell attachment to the surface. Two transfection mixture tubes, one containing 6 μl Cellfectin II (Invitrogen, #10362100) in 250 μl Sf-900 II SFM, and another containing 2 μg bacmid DNA in 250 μl SF-900 II SFM, were prepared. After 5 min, the two mixtures were combined and incubated for 20 min at room temperature. The transfection mix was then added dropwise to the Sf9 cells. Cells were incubated at 27°C for 72 hours, and the GFP expression was monitored by fluorescence microscopy. The supernatant containing the P1 virus was collected, filtered through a 0.2 μM filter, supplemented with 2% FBS, and stored at 4°C. The P1 virus was amplified by infecting 50 ml of Sf9 cells to generate P2, which was subsequently used to prepare the P3 virus. The P3 viral titer was determined using Sf9 Easy Titer cells, as described previously^90^.

To transduce the Expi293 cells, cells were grown in Expi293 expression medium at 37°C in the presence of 8% CO_2_ and 80% humidity until a cell density of 2.5 × 10^6^ cells/ml. The cells were transduced with TrkA-GFP-FLAG baculovirus (P3) at a multiplicity of infection (MOI) of 1 and incubated at 37°C. After the transduction, the culture was incubated in an incubator under conditions mentioned above.

### Lentivirus production

For lentiviral production, HEK293FT cells were seeded in 10 or 15 cm dishes (Corning, #430167 or Celltreat, #229651) and transfected at ∼80% confluency using PEI as described above. The 2^nd^ or 3^rd^ generation lentiviral plasmids are transfected along with 2^nd^ or 3^rd^ generation packaging systems, respectively, according to manufacturer’s protocol (2^nd^ generation: pEXQV, pMD2.G; 3^rd^ generation: pMD2.G, pRSV-Rev, and pMDLg/pRRE). These DNA and PEI were diluted separately in Opti-MEM and incubated for 5 minutes, following which they were combined and incubated for 20 minutes, before being added dropwise to the cells. After 10–12 hours, the medium was replaced with fresh DMEM + 10% FBS (without antibiotics). Virus-containing supernatant was harvested at 24 and 48 hours, clarified by centrifugation, and concentrated with a Lenti-X concentrator (TAKARA, #631231) following manufacturer’s instructions. Concentrated lentivirus was then centrifuged at 1500g for 45 min at 4°C. The pellet obtained was resuspended in an appropriate volume of DMEM + 10% FBS medium. The concentrated virus was aliquoted and stored at -80°C until further use.

### Lentiviral Transduction

SHSY5Y cells were seeded a day before transduction to reach 70–80% confluency at the time of infection. For each 10 cm plate, transduction was carried out in 7 mL of Advanced DMEM/F12 medium supplemented with 10% FBS, 1 mM L-glutamine, and 1 mM AA. Lentivirus (50× concentrated, 125 µL) and polybrene (6 µg/mL, EMD Millipore, #TR-1003-G) were mixed with the media and carefully added to the cells. After 12 h, the viral media were replaced with fresh growth medium. Transduction efficiency in SHSY5Y cells was monitored over 60–72 hours by assessing expression of GFP. For HEK293FT cells, transduction was performed at 70–80% confluency using the same procedure. A total of 50 µL concentrated virus was added to 7 mL medium containing 6 µg/mL polybrene. After 12 hours, the medium was replaced with fresh DMEM supplemented with 10% FBS and 1 mM AA. Fluorescent tag expression (GFP or mCherry) was monitored over 24 hours, and cells were harvested for flow cytometry-based Fluorescence-Activated Cell Sorting (FACS) once desired levels of expression were detected.

## Stable cell line generation and maintenance

To generate SHSY5Y or HEK293 stable cell lines expressing different versions of TrkA (**Supplementary Table 3**), cells were transduced with appropriate lentivirus carrying the TrkA construct of interest. Cells were transduced at ∼80% confluency with lentivirus as above. After 12 hours, the viral medium was replaced with fresh media, and cells were cultured for an additional 24-60 hours. Transduced cells expressing the fluorescent reporter (e.g., TrkA-GFP-HIS/FLAG) were sorted using a SONY-MA900 cell sorter. For Fluorescence-Activated Cell Sorting (FACS), the cells were resuspended in FACS buffer (DPBS + 10% FBS) at a density of 1-2 million/mL, filtered through a 40 μm strainer (Falcon, #352235), and sorted based on fluorescence intensity into low, medium, and high bins. Polyclonal populations were expanded, re-sorted if needed, and validated for stable expression by fluorescence microscopy and immunoblotting.

To generate a JGCaMP8m and TrkA co-expressing stable cell line, SHSY5Y-TrkA stable cells (Kerafast, #ECP006) were transduced at ∼80% confluency with 125 µL of 50× concentrated jGCaMP8m lentivirus in 7 mL of culture medium supplemented with 6 µg/mL polybrene. Virus-containing medium was removed after 12 hours, and cells were further incubated for a total of 72 hours post-transduction before FACS based on jGCaMP8 basal fluorescence into low, medium, and high expressing populations. Sorted cells were pelleted, resuspended in 1 mL of complete medium without G418, and seeded into 6-well plates with 2 mL of RPMI media per well. Cells were allowed to attach and recover. The media was changed every 60 hours. At 96 hours post-sorting, G418 was added at a final concentration of 0.3 mg/mL to maintain the integrated TrkA in the cells and the antibiotic was subsequently maintained at this concentration.

To generate a TrkA-GFP-His and hTRPV1 co-expressing HEK293 stable cell line, we transduced cells with an EF1A-driven hTRPV1 lentiviral vector also simultaneously co-expressing a fluorescent marker mCherry driven by a CMV promoter from a separate cassette (mCherry/Puro-EF1A>hTRPV1). Following transduction, GFP and mCherry fluorescence confirmed expression of TrkA-GFP and hTRPV1, respectively. Cells were prepared as described above and sorted by flow cytometry for dual GFP and mCherry-positive populations. The sorted cells were expanded and used for electrophysiology experiments. All cell lines used in this study are summarized in **Supplementary Table 3**.

## Cellular treatment with NGF variants

For time-course experiments, TrkA-GFP-His-expressing SHSY5Y cells were serum starved for ∼14 hours before treatment with NGF or NGF^painless^ (50 ng/mL) for the indicated time points (2.5, 5, 15, 30, or 60 min) at 37°C. Serum-starved mock-treated cells (no NGF) were used as controls. For endpoint experiments, cells were serum starved as above, and either left untreated or stimulated with NGF or NGF^painless^ (50 ng/mL) for 15 min at 37°C_._ For NGF concentration titration experiments, cells were serum starved as above and treated with varying NGF concentrations (0, 10, 20, 50, 100, or 200 ng/mL) for 15 min at 37°C. At the end of each treatment, culture medium was aspirated, washed with ice-cold 1× DPBS, harvested with 0.05% trypsin, spun down at 1000g for 3 min at 4°C, and flash frozen until further use. NGF was a gift from Genentech, USA, and NGF^painless^ was obtained from the European Brain Research Institute.

## Immunoblotting, in-gel fluorescence, and Coomassie Brilliant Blue staining

### Western blot sample preparation

For Western blot, cell pellets were lysed in 300 µL of 1× Cell Lysis Buffer (Cell Signaling Technology, #9803) supplemented with protease inhibitor cocktail (Thermo Fisher Scientific, #A32953), phosphatase inhibitors cocktail 2 and 3 (Sigma-Aldrich, PIC2; #P5726, PIC3; #P0044 ), 2 mM sodium orthovanadate, 50 mM sodium fluoride and 1 mM PMSF. Lysates were incubated on ice for 30 min, centrifuged at 10,000g for 10 min at 4°C, and clarified supernatants were collected. GFP fluorescence was measured using a ChemiDoc system to normalize protein input. About equal amounts of protein were mixed with 4× Laemmli buffer (Bio-Rad, #1610747) containing 50 mM DTT and heat-denatured at 70°C for 7 min.

### Western blotting

Proteins were resolved on 4–20% or 8-16% Mini-PROTEAN TGX Stain-Free gels (Bio-Rad, #4568096, #4561106) and transferred to PVDF membranes (Bio-Rad, #1620177) using either a semi-dry or wet transfer system. Membranes were blocked with 4% Bovine Serum Albumin (BSA, Sigma-Aldrich, #A9647) in TBST (Tris-Buffer saline with 0.4% Triton X-100) for 1 h at room temperature, incubated overnight at 4 °C with primary antibodies. The list of antibodies used for immunoblotting are summarized in **Supplementary Table 5**. Primary antibodies were detected with species-specific secondary antibodies linked to HRP for chemiluminescence detection. GAPDH and calnexin were used as loading controls. Blots were developed using Clarity Western ECL substrate (Bio-Rad, #1705000).

### Immunoblot and SDS-gel imaging and analyses

Blots of target bands were acquired using either the chemiluminescence or fluorescence channels (based on the secondary antibody used) on a ChemiDoc MP system (Bio-Rad). Protein molecular weight ladders were imaged using the colorimetric channel, and these images were merged with target band images to detect molecular weights of observed bands. For in-gel GFP fluorescence, following SDS-PAGE, the gels were washed 2× with about 50 mL of ddH_2_O before laying the gels directly on the appropriate imaging tray and acquiring images using the Alexa488 detection channel in the ChemiDoc MP system (Bio-Rad). Protein molecular weight ladders were imaged using the Coomassie blue channel, and these images were merged with Alexa-488 images for visualization.

Immunoblots and gels were quantified using the Gel Analyzer function in Fiji^61^. Briefly, bands were selected using a rectangular ROI, and raw band intensity was calculated for each selected band by integrating the area under the curve (AUC) of the observed intensity distribution produced by the Gel Analyzer function. For proteins of interest, band intensity values were normalized to those of loading control proteins (calnexin and GAPDH). Phosphorylated proteins were normalized to the signal of loading controls and to total protein levels.

## Calcium signaling assay

For calcium signaling assays, the TrkA- and jGCaMP8m-expressing SHSY5Y stable line was used. Cells were grown ∼90% confluency and transferred to PDL-coated MatTek dishes (#P35GF-1.5-14-C) at a density of 0.39 million cells per dish in 2 mL of pre-warmed serum-free Advanced DMEM/F-12 supplemented with 0.3 mg/mL G418, and incubated at 37°C, 5% CO₂ for 14 h. Before imaging, the media was replaced with serum-free, phenol red–free RPMI medium containing 0.8 mM EGTA. We included EGTA in the imaging media to ensure that only Ca²⁺ released from intracellular stores was monitored. Cells were stimulated with NGF or NGF^painless^ at ∼1.5 min into imaging at a final concentration of 100 ng/mL. For the control samples, an equivalent volume of 1× DPBS was added at the same time frame during imaging. Time-lapse images were acquired to monitor the increase in intracellular calcium levels in response to NGF variants.

Time-lapse calcium imaging was performed on a Nikon Ti2 widefield fluorescence microscope with an S Plan Fluor ELWD 40×/0.60 Ph2 objective and the Perfect Focus System. Illumination came from a Lumencor SOLA II LED light engine passed through a Chroma 49002 ET-GFP filter set (ET470/40x excitation; ET525/50m emission). Images were acquired on a Photometrics Prime 95B sCMOS camera in 16-bit, 1×1 binning at 1024

× 1024 pixels (0.2788 µm/pixel; field of view 285.5 µm × 285.5 µm). Each frame used a 200 ms exposure, and excitation power at the sample plane was 915 µW. Each series comprises 40 frames at ∼30 s intervals (total duration ∼20 min). Time-lapse images were acquired in a live cell incubation chamber that maintained a constant temperature of 37°C and 5% CO_2_. Time-lapse image stacks were automatically processed in Fiji^61^ (ImageJ 2.16.0) with a custom macro. Briefly, each stack was first preprocessed by correcting XY drift with Fast4DReg^91^, then subjected to background subtraction using the rolling-ball method (radius 50 pixel radius). For segmentation, the slice with the highest mean intensity in each stack was used to generate cell masks by threshold-based segmentation, and only ROIs with an area ≥ 25 px² were used for analysis. The resulting ROIs were applied to the full time series so that the same cells were measured across frames, yielding per-cell integrated-density (IntDen) traces. For each cell, ΔF/F₀ was computed by normalizing the IntDen trace to its own pre-stimulation baseline F₀, and the resulting ΔF/F₀ time courses were used to quantify calcium transients.

## Single particle tracking in live cells and data analysis

### Sample chamber preparation

Single particle tracking samples were prepared in 35 mm glass-bottom MatTek dishes (No. 1.5, uncoated, gamma-irradiated, #P35G-1.5-14-C) after cleaning the dishes by incubating with 1 mL freshly-made 1 M KOH (Sigma-Aldrich, #221473) solution for 15 minutes at room temperature, followed by aspiration, and 3× wash with 2 mL of MilliQ H_2_O. During the third wash, the entire dish was thoroughly washed by vigorously pipetting MilliQ H_2_O before removal by aspiration. The MatTek dish was then cleaned once more by treatment with 2 mL of 100% ethanol for 15 minutes at room temperature. The dish was again washed thrice with 2 mL MilliQ H_2_O as above. These MatTek dishes were coated with poly-L-lysine by incubating the glass coverslip with 400 μL of 0.01% poly-L-lysine solution (EMD Millipore, #A-005-C) for 15 minutes at room temperature. The dish was washed as described above and left in 2 mL of sterile DPBS until cell seeding.

### Cell seeding, starvation, and labelling

Sorted cells stably expressing ALFA-TrkA-GFP-FLAG were grown to 90-95% confluency in a 10 cm dish and split using trypsinization as described previously. 1.5 million re-suspended cells were seeded in each MatTek dish, supplemented with 2 mL of SHSY5Y media (supplemented Advanced DMEM/F12 media as described above). These dishes were incubated at 37°C to allow the cells to attach to the poly-L-lysine-coated surface for

7-8 hours. After visual confirmation of cell attachment using microscopy, the conditioned media was gently aspirated and replaced with starvation media (SHSY5Y media without FBS). Cells were returned to the 37°C incubator and starved for ∼15 hours.

Starved cells were then labelled with a FluoTag X2 anti-ALFA nanobody conjugated with Alexa-647 dye (Nanotag, #N1502-AF647-L). For labelling, the nanobody was serially diluted to a final concentration of 30 pM in pre-warmed labelling media, composed of phenol-free DMEM/F12 media (Gibco, #15630080) supplemented with L-glutamine, antimycotic-antibiotic, 25 mM HEPES, and 5% BSA. The starvation media was aspirated and replaced with 2 mL of 30 pM nanobody labelling solution, and the dish was returned to the 37°C incubator for 15 minutes. The labelling solution was aspirated from the dish, and cells were washed with 2 mL of DPBS before addition of 1.8 mL of imaging media, composed of phenol-free DMEM/F12 media supplemented with L-glutamine, antimycotic and antibiotic, 25 mM HEPES, to the cells. At this stage, cells were ready for TIRF imaging and immediately transported to the microscope.

### Single particle tracking data acquisition

Cells were imaged using a Nikon Ti2-E inverted microscope, first by mounting MatTek plates on a motorized stage and engaging a Perfect Focus system for stable focusing throughout image acquisition. Cells were illuminated using a Nikon LU-N4 unit with 488 nm, 561 nm, and 640 nm solid state lasers, manipulated by a TIRF illuminator and imaged using a Nikon 100 x 1.49 oil immersion TIRF objective employed in 1.5x magnification mode. These lasers were controlled using an acousto-optic tunable filter (AOTF), and a Quad TIRF filter set from Chroma Technology Corp. (405/488/561/638 nm) was employed with additional emission filters for each laser channel: 700/75 m for the 488 nm channel, 525/50 m for the 561 nm channel, and 600/50 m for the 640 nm channel. Images were recorded using an Andor iXon Life 897 EMCCD camera, operated with 16-bit, 1x1 binning at 512 x 512 pixels (0.256 um/pixel), and a field-of-view of 54.05 x 54.05 um. All imaging was operated with the use of NIS elements software.

The workflow for single particle tracking data acquisition consisted of the following procedures. Flat, well-adhered cells expressing ALFA-TrkA-GFP (determined by imaging in the 488 channel) were identified and their locations indexed using NIS elements image acquisition mode. Single frame 488 nm images were captured for each cell to document their TrkA expression levels (5.9 mW laser power, 30 ms exposure, EM gain 58). Anti-ALFA-Nanobody labels, conjugated to Alexa-647 dye, were photoblued to 561 nm species by illuminating each cell with the 640 nm laser (3.7 mW power, EM gain 300) in stream acquisition mode for 500 frames using a 30 ms exposure time, as previously reported^70^. After photoblueing, a subset of the nanobody-alexa-647 is converted to 561 nm emitting species. The short-lived species among these 561 nm photoblued labels were bleached out of the sample by pre-illuminating each cell with the 561 nm laser (6.3 mW power, EM gain 300) for 500 frames using a 30 ms exposure time. After this photobleaching step, the photoblued species that remained in the selected cells were of a photostable, long-lived character suitable for single particle tracking acquisition, as previously shown^70^.

To record single particle tracking data, cells were illuminated with the 561 nm laser (6.3 mW laser power, EM gain 300) for 1500 frames using a 30 ms exposure time. Each cell was imaged before the addition of NGF to establish baseline TrkA tracking data. After this initial acquisition, the cells were treated with NGF at a final concentration of 100 ng/mL, while still mounted on the scope. The NGF was allowed to diffuse throughout the sample for 15 minutes at room temperature, before “post-NGF treatment” TrkA tracking datasets were collected for each cell for the next ∼15 min (until a total of ∼30 minutes having elapsed after NGF treatment). Data from a total of ∼55 cells were collected across multiple plates and over three biological replicates before and after treatments with NGF or NGF^painless^.

Image stacks were analyzed in MATLAB R2024a with custom routines built on the u-track 2.5 framework as previously described^70,92^. Each 16-bit TIRF frame was first divided by the camera full-scale value and filtered with a Gaussian kernel whose standard deviation matched the experimentally measured point-spread function 𝜎_!"#_. Candidate particles were identified as local maxima whose amplitudes exceeded the masked local background by a one-tailed normal test. Every candidate was then refined by non-linear least-squares fitting to an isotropic two-dimensional Gaussian plus constant background,

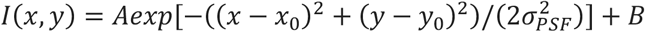

yielding sub-pixel coordinates (𝑥_0_, 𝑦_0_) together with localisation errors propagated from the fit covariance. Detection was executed in parallel, with each CPU worker processing an independent frame block; the resulting particle lists were merged into a single structure for downstream analysis. Inter-frame association relied on the native u-track assignment solver. Coordinates were converted to micrometers using the calibrated pixel size, 𝑣*_px_*. For every trajectory 𝑖 the mean-squared displacement was calculated as

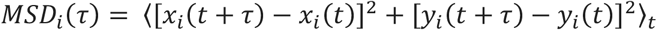

### Rolling diffusion coefficient determination and trajectory major and minor axis extraction

An eleven-frame centered window were slid along each track; within each window successive displacements were different, giving an effective lag time Δ𝑡*_eff_*. The local diffusion coefficient followed the free-Brownian relation

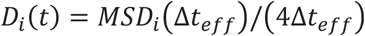

Time-averaging 𝐷*_t_*(𝑡) provided one diffusion value per trajectory, and the ensemble mean of those values defined the cellular diffusion coefficient. Additional trajectory descriptors were derived to characterize motility geometry and persistence. To quantify track shape, principal-component analysis of the positional cloud produced eigenvalues 𝛾_1,_ ≥ 𝛾_2_ and the full major axes of the trajectory ellipse were calculated as 2_√_𝛾_,_.

## Native-nanoBleach to determine oligomeric distribution

### Solubilization of TrkA-NGF complex in styrene maleic acid (SMA)-derived native nanodiscs

Sorted HEK293FT cells stably expressing TrkA-GFP were grown in 15 cm plates in the presence of TrkA inhibitor at 5 μM. Actively growing cells at 80% confluency were serum-starved for ∼14 hours. Individual plates were treated with 100 ng/ml NGF or NGF^painless^ for 4 min at 37 °C. The cells were quickly harvested using 0.05% trypsin, washed with 1× cold DPBS, and stored at -80 °C until use. The solubilization of TrkA:NGF complex in native nanodiscs using SMA is done as previously described^62^. Briefly, harvested cells were lysed by resuspending in lysis buffer (**Supplementary Table 11**) and lysed using a nitrogen cavitation chamber pressurized to 1000 PSI for 3 min, removing the cell debris by soft spin at 3000g for 7 min. The supernatant was collected and subjected to ultracentrifugation at 200,000g for 45 min to pellet membranes. The membrane pellet was then resuspended in membrane resuspension buffer (**Supplementary Table 11**) using a Dounce homogenizer. The resuspended membrane was solubilized using 2% SMA (Orbiscope, #SMALP 200) for 90 minutes at 4°C. The soluble material was then subjected to ultracentrifugation again to pellet any insoluble materials and isolate the soluble fraction in the supernatant. This soluble fraction contained native nanodiscs derived from the whole membrane fraction of the sample.

The soluble native nanodisc sample were enriched and analyzed by fluorescence-detected size-exclusion chromatography (FSEC) on a Superose 6 10/300 GL column (Cytiva) using an ÄKTA pure FPLC system coupled in-line to a Shimadzu RF-20AXS fluorescence detector (GFP monitored at 488 nm). FSEC traces resolved native nanodisc-encapsulated TrkA-GFP:ligand complex from soluble aggregates (void volume) and from smaller proteolysis/degradation products eluting at later volumes. Non-void fractions corresponding to the full-length TrkA-GFP:NGF were verified by SDS–PAGE by in-gel GFP fluorescence (samples prepared without heating or reducing agent), confirming the molecular weight of the nanodisc-encapsulated, TrkA-GFP-tagged protein^62^. GFP-positive, non-void fractions were chosen and subjected to Native-nanoBleach^62^. This same protocol can be used for other types of cells as well.

### Sample preparation, data acquisition and analysis

Sample preparation, data acquisition, and processing were performed as described before^62^. Briefly, glass coverslips (ibidi, #80608) were sequentially cleaned with detergent, isopropanol/water, and water before being plasma cleaned and affixed with microfluidic flow chambers (Ibidi, #80608-90). In each microfluidic well, glass was passivated with PLL-PEG/PEG-biotin, and then functionalized for GFP-nanodisc capture by coating first with streptavidin and then biotinylated GFP-nanobody. FSEC-eluted nanodisc samples were incubated in functionalized wells for 5 minutes at room temperature, allowing for immobilization of nanodisc-encapsulated, GFP-tagged proteins directly on the glass. The dilution of streptavidin, GFP-nanobody, and FSEC-eluted nanodisc sample were optimized to immobilize GFP-tagged nanodiscs at single-particle density as previously established^62^. After sample incubation, each well was washed 3× with 1 mL DPBS.

Single-particle TIRF images were acquired on a Nikon Eclipse Ti2-E inverted microscope equipped with Nikon 100 × 1.49 numerical aperture oil-immersion TIRF objective, a TIRF illuminator, a Perfect Focus system, and a motorized stage. Images were recorded using an Andor iXon electron-multiplying charge-coupled device camera. Tricolor images were recorded at 20 disparate locations in each well by illuminating with 488 nm, 561 nm, and 640 nm lasers (5.9 mW, 9 mW, 11.2 mW, respectively, 80 ms exposure for each) coupled with their respective filter sets. Tricolor images confirmed the single-molecule resolution of the on-glass nanodisc capture and allowed visualization of any background signal in the 561 nm and 640 nm channels as quality control. Photobleaching experiments were recorded in stream acquisition mode (1,500 frames at ∼9.3 fps, 80 ms exposure) under continuous 488 nm illumination at 1.8 mW and 2.7 mW for each sample. Single particles were localized with TrackMate (Fiji/ImageJ)^93^, and fluorescence traces were extracted using custom MATLAB code^62^ to determine discrete GFP photobleaching steps. Traces showing 1–4 clear steps were scored, and distributions from ∼1,000–1,500 particles per replicate were used to calculate monomer, dimer, trimer, and tetramer fractions, correcting for 70% GFP maturation efficiency, over 4 biological replicates, as described previously^62^.

## Cryo-EM studies of TrkA:NGF and TrkA:NGF^painless^ complex in reconstituted native-like lipid nanodiscs

### TrkA protein expression

Expi293 cells were grown in Expi293 expression medium at 37°C in the presence of 8% CO_2_ and 80% humidity until a cell density of 2.5 × 10^6^ cells/ml. The cells were transduced with TrkA-GFP-FLAG baculovirus at a multiplicity of infection (MOI) of 1 and incubated at 37°C. TrkA inhibitor (Selleckchem, #GW441756; **Supplementary Table 8**) was added to transduced cells at a concentration of 5 µM and maintained throughout expression and purification. After 10 hours, transduced cells were supplemented with 10 mM Sodium butyrate (Sigma-Aldrich, #303410), and the temperature was reduced to 30°C. After 48 hours, cells were treated with 4 μg/ml NGF (or NGF^painless^) for 15 min at 37°C. Cells were harvested by centrifugation, flash-frozen, and stored at -80°C until use.

### Membrane solubilization and protein purification

Harvested cells were resuspended in cell lysis buffer (25 mM Tris pH 8.0, 150 mM KCl, 2 μM TrkA inhibitor (Selleckchem, #GW441756), protease inhibitor cocktail tablet (Thermo Fisher Scientific, #A32953), 2 mM sodium orthovanadate (EMD Millipore, #567540), 50 mM Sodium fluoride (Sigma-Aldrich, #201154), and 2 mM PMSF (Sigma-Aldrich, #52332). The resuspended cells were lysed using a nitrogen cavitation chamber at 1000 psi for 7 minutes. The cellular debris were cleared by a soft spin at 3000g for 7 min. The supernatant from soft spin was subjected to hard spin at 200,000g for 45 min to isolate the total membrane using a Ti70 rotor in an ultracentrifuge (Beckman Coulter). The isolated membrane was resuspended in membrane resuspension buffer (25 mM Tris, pH 8.0, 150 mM NaCl, 5% glycerol, 2 μM TrkA inhibitor (GW441756), and supplemented with protease inhibitors) using a Dounce homogenizer. To initiate the solubilization, 1% DMNG (Anatrace, #NG322) was added, and the mixture was nutated for 90 min at 4 °C to solubilize and extract the TrkA from the membrane.

The solubilized mixture was centrifuged at 200,000*g* for 45 min, and the supernatant was collected and incubated with pre-equilibrated anti-FLAG-M2 affinity gel (Sigma Aldrich, #A2220-5ML, **Supplementary Table 9**) for 60 min at 4°C with gentle mixing. The suspension was then passed through a gravity column. Beads were washed with 30 mL of membrane resuspension buffer with 0.1% DMNG, and protein was eluted with elution buffer containing 0.2 mg/ml FLAG peptide in elution buffer supplemented with 0.1% DMNG. NGF was maintained at a concentration of 1 μg/ml throughout the purification process, including cell lysis, solubilization, FLAG enrichment, and elution. The eluted protein was concentrated using a Viva Spin Concentrator (100 kDa cutoff) and used for nanodisc reconstitution. All purification buffers are summarized in **Supplementary Table 11**.

### Reconstitution of TrkA:NGF/NGF^painless^ complexes into membrane scaffold protein (MSP) lipid nanodiscs

The TrkA:NGF protein complex (WT or painless, TrkA:NGF^WT/painless^) was reconstituted into nanodiscs as described before^77^. Briefly, a 50 mM stock of lipids was made by weighing 38 mg of soy phospholipids (Millipore Sigma, #11145), which were dissolved in chloroform and completely dried under a nitrogen stream. The dried lipid film was resuspended in 1× TBS buffer (25 mM Tris, pH 7.5, 200 mM NaCl). The suspension was subjected to five freeze–thaw cycles (freezing in liquid nitrogen followed by thawing in a bath sonicator). The resulting homogeneous lipid stock (50 mM) was either used immediately or aliquoted and stored at −80 °C. The membrane scaffold protein MSPE3D1 was expressed and purified from *E. coli* following the reported protocol^94^. The purified TrkA:NGF or TrkA:NGF^painless^ complex was supplemented with an excess of corresponding NGF variants (∼100 μg/ml or ∼4 μM; wild-type or painless variant) and incubated on ice for 1 h before nanodisc assembly. Protein, lipid, and MSP were mixed at a molar ratio of 1:6:600 and incubated on ice for 1 hour. To initiate nanodisc formation, pre-activated Bio-Beads SM-2 (Bio-Rad, #1528920) were added to remove detergent, and the mixture was left for ∼12 hours at 4°C with end-to-end rotation. The Bio-Beads were then replaced with fresh beads at 4°C for 2 hours. This step was repeated once more with new Bio-Beads for an additional 2 hours. After reconstitution, the mixture was centrifuged at 10,000*g* for 10 minutes at 4°C to remove any aggregates. The clarified sample was purified by size exclusion chromatography (SEC), and the fraction corresponding to the dimeric TrkA:NGF complex was collected. This fraction was concentrated, analyzed by SDS-PAGE and FSEC, and used for cryo-EM grid preparation.

### Vitrification and cryo-EM data collection

Holey-gold 300 mesh R1.2/1.3 µm grids (Ultra Au Foil R1.2/1.3 300 mesh, EMS, #Q350AR13A) were glow discharged (SCD005 Sputter Coater, Bal-Tec) for 1 min at 15 mA current and ∼0.1 mBar air pressure. Glow-discharged grids were immediately used for sample vitrification. 3µL of purified TrkA:NGF at 0.22 mg/mL and TrkA:NGF^painless^ at 0.23 mg/mL were applied to the grid and blotted using a Vitrobot Mark IV (Thermo Fisher Scientific) with the following settings: *Blot time* – 2.5 seconds; *blot force –* 0; *temperature* – 12 ^°^C; *humidity* – 100%. Movies were recorded using a K3 (Gatan Inc.) direct electron detector in counting mode at a nominal magnification of 105,000× resulting in calibrated pixel size of 0.832Å per pixel using Serial EM^95^ on a Titan Krios G4i transmission electron microscope (Thermo Fisher Scientific) operating at 300 kV equipped with a bioQuantum energy filter with slit width set to 20 eV. Exposures were recorded for 2 seconds and were dose fractionated into 50 frames, yielding a total dose of ∼50 e^-^ Å^-2^, and a defocus range of -2.5 to -1.0 µm.

### Cryo-EM data processing

All image processing was carried out using CryoSPARC v4.2.1^96^. Briefly, motion correction was carried out on the raw movies, and the contrast transfer function (CTF) was estimated using patch CTF correction on the dose-weighted motion-corrected averages. A total of 6,834 and 11,831 micrographs were selected for further processing based on a 7 Å CTF cutoff for TrkA:NGF and TrkA:NGF^painless^ datasets, respectively. Particles were initially picked using the blob-picker to generate 2D templates for template picking. Template-picking was then performed on the entire dataset, and particles were extracted with a box size of 400 pixels, Fourier-cropped to 200 pixels, and subjected to iterative 2D classification to discard obvious junk particles. Selected particles were used for iterative rounds of *ab initio* to remove junk and monomeric TrkA particles. Dimeric TrkA:NGF particles were then re-extracted without binning and subjected to homogeneous and non-uniform refinements. For further refinement, CryoSPARC was updated to v4.6.2 to implement reference-based motion correction on select particles. Non-uniform refinement was then performed on the polished particles, followed by global and local CTF refinements and another round of non-uniform refinement. Local refinement jobs were then conducted using masks that encompass the entire ECD (4.02Å and 3.13Å) or only the Ig2 domain of TrkA-ECD and NGF homodimer (3.79Å and 2.87Å), for TrkA:NGF and TrkA:NGF^painless^ datasets, respectively.

### Cryo-EM model building and validation

Crystal structure of the ECD of TrkA in complex with NGF (PDB: 2IFG)^42^ was used as a template for model building. Appropriate mutations (P61S and R100E) were introduced in the NGF for the NGF^painless^. The models were fit into the experimental Coulomb potential map using molecular dynamics flexible fitting (MDFF)^97,98^. Atomic positioning derived from MDFF calculations was first assessed and improved using real-space refinement in Phenix 1.21.1 and then final manual fine tuning was done using Coot v0.9.8.94^99^.

## Electrophysiology studies

HEK293T cells stably expressing TrkA-GFP and TRPV1 were serum-starved in DMEM for ∼6-8 hours before experiments. Cells were detached using Detachin™ cell detachment solution (Avantor, #MSPP-T100110) and resuspended in serum-free DMEM at a density of 1×10^7^ cells/mL. The cell suspension was then loaded into an 8-well QPlate, and whole-cell patch-clamp recordings were performed using the QPatch Compact automated patch-clamp system (Sophion Bioscience). The extracellular solution (EC) contained 140 mM NaCl, 1.8 mM CaCl₂, 1 mM MgCl₂, 4 mM KCl, and 10 mM HEPES; pH was adjusted to 7.4 with NaOH, and osmolarity was adjusted to 310 mOsm with D-mannitol. The intracellular solution (IC) contained 80 mM KF, 55 mM KCl, 1.6 mM MgCl₂, 2 mM EGTA, 2.5 mM Na₂ATP, 0.2 mM Li₂GTP, and 10 mM HEPES; pH was adjusted to 7.3 with NaOH, and osmolarity was adjusted to 315 mOsm with D-mannitol.

After establishing the whole-cell configuration, the membrane potential was held at -60 mV, and current responses were recorded. 5 μL of 10 μM capsaicin prepared in EC was applied as a 1-second pulse at 30 second intervals until stable current responses were obtained. After stabilization, 5 instances of capsaicin-evoked currents were recorded. Subsequently, 100 ng/mL NGF or NGF^painless^ was perfused at a final volume of 10 μL and incubated at 25°C for 15 minutes. Following this incubation, capsaicin-evoked currents were again recorded 5 times from the same cells using identical capsaicin stimulation protocol as above.

## Native mass spectrometry

NGF and NGF^painless^ were buffer exchanged into 200 mM NH4Ac with Zeba spin desalting columns (Thermo Fisher Scientific, #89877). The protein concentration was kept at 10 µM for optimal signal. The sample was ionized via nano electrospray ionization with the nano-emitter capillaries formed by pulling borosilicate glass capillaries (outer diameter: 1.5mm, inner diameter: 1.1 mm, length: 7.5cm; Sutter Instruments) using a Flaming/Brown micropipette puller (Sutter Instruments, Model P-1000) equipped with 3.0 × 3.0 mm^2^ box filament (Sutter Instruments). All nMS and nTD-MS data were acquired by Q Exactive UHMR (Thermo Fisher Scientific), which was modified with ExD TQ-160 cell (e-MSion, Agilent). Thermo Tune software (Q Exactive UHMR 2.13 Build 3159) was used to control the instrument, and the data was analyzed using FreeStyle 1.7 (Thermo Fisher Scientific), ExDViewer Version 4.5.6 (MS2, e-MSion, Agilent).

Once the broadband spectra were collected for NGF and NGF^painless^, the 11+ charge states of the dimers were isolated and then subjected to activation via higher-energy collisional dissociation (HCD) at 30V and 17V, respectively. The energy level was optimized to induce monomer ejection without fragmentation. The resulting dimer and monomer spectra were manually deconvoluted using an in-house script. After nMS and nTD-MS analysis, the NGF and NGF^painless^ aliquots were subjected to stress through freeze-thaw cycles and storage at 4°C for 5 days. These proteins were then re-analyzed with the same nMS condition as above.

## APBS calculations

Electrostatic surface potentials were computed with the APBS plugin in PyMOL v3.1.6.1. Structures were protonated and charge-assigned using PDB2PQR. The Poisson–Boltzmann equation was solved under aqueous conditions with default grid settings. The resulting potential maps were projected onto the solvent-accessible molecular surface and visualized using PyMOL with a color scale ranging from -5 to +5.

