## Supplementary Figures for "A painless nerve growth factor variant uncouples nociceptive and neurotrophic TrkA signaling"

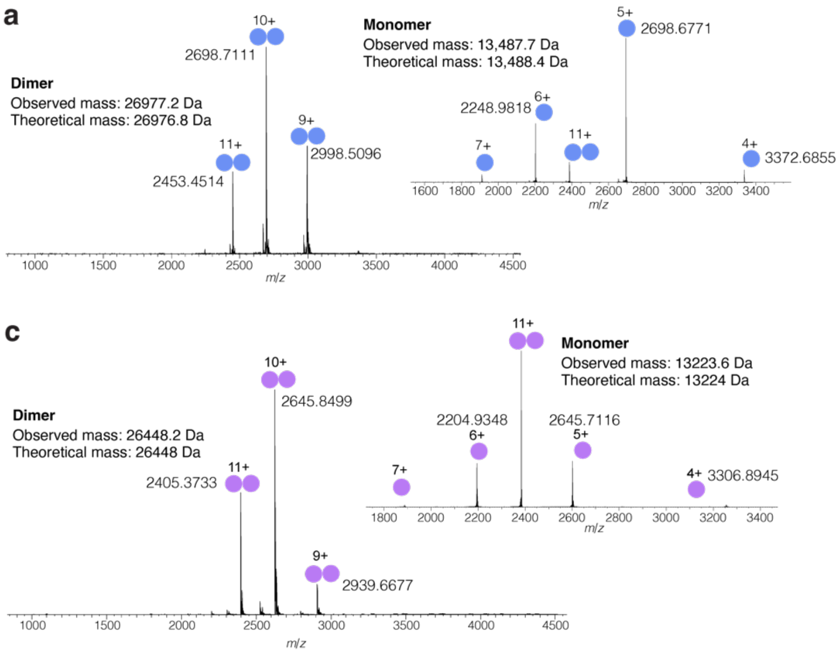


**Supplementary Figure 1**: **Native mass spectrometry analysis (nMS) of NGF and NGF$^{painless}$**. **a.** nMS of NGF clearly showed the dimeric status of this protein. **Inset.** The 11+ charge state of the dimer was isolated and gently activated to induce monomer ejection. The majority of the population splits into a 5+/6+ monomer pair. The observed dimeric and monomeric masses are in agreement with the theoretical molecular weights, indicating that the six disulfide bonds reported in NGF are fully intact. **b.** The NGF dimer was freeze-thawed and stored for ~5 days at 4 °C and then subjected to nMS analysis again. The resulting spectrum and the corresponding observed mass are nearly identical to those shown in (a), indicating high innate stability of the dimer. **c.** nMS analysis of NGF$^{painless}$ clearly showed the dimeric state of this protein. **Inset.** The 11+ charge state of the dimer was isolated and gently activated to induce monomer ejection. Again, the majority of the population splits into a 5+/6+ monomer pair. **d.** The NGF$^{painless}$ dimer was freeze-thawed and stored for ~5 days at 4 °C and then re-analyzed via nMS. The resulting spectrum and the corresponding observed mass are nearly identical to those shown in (c), again indicating high innate stability of the dimeric state of the mutant NGF.

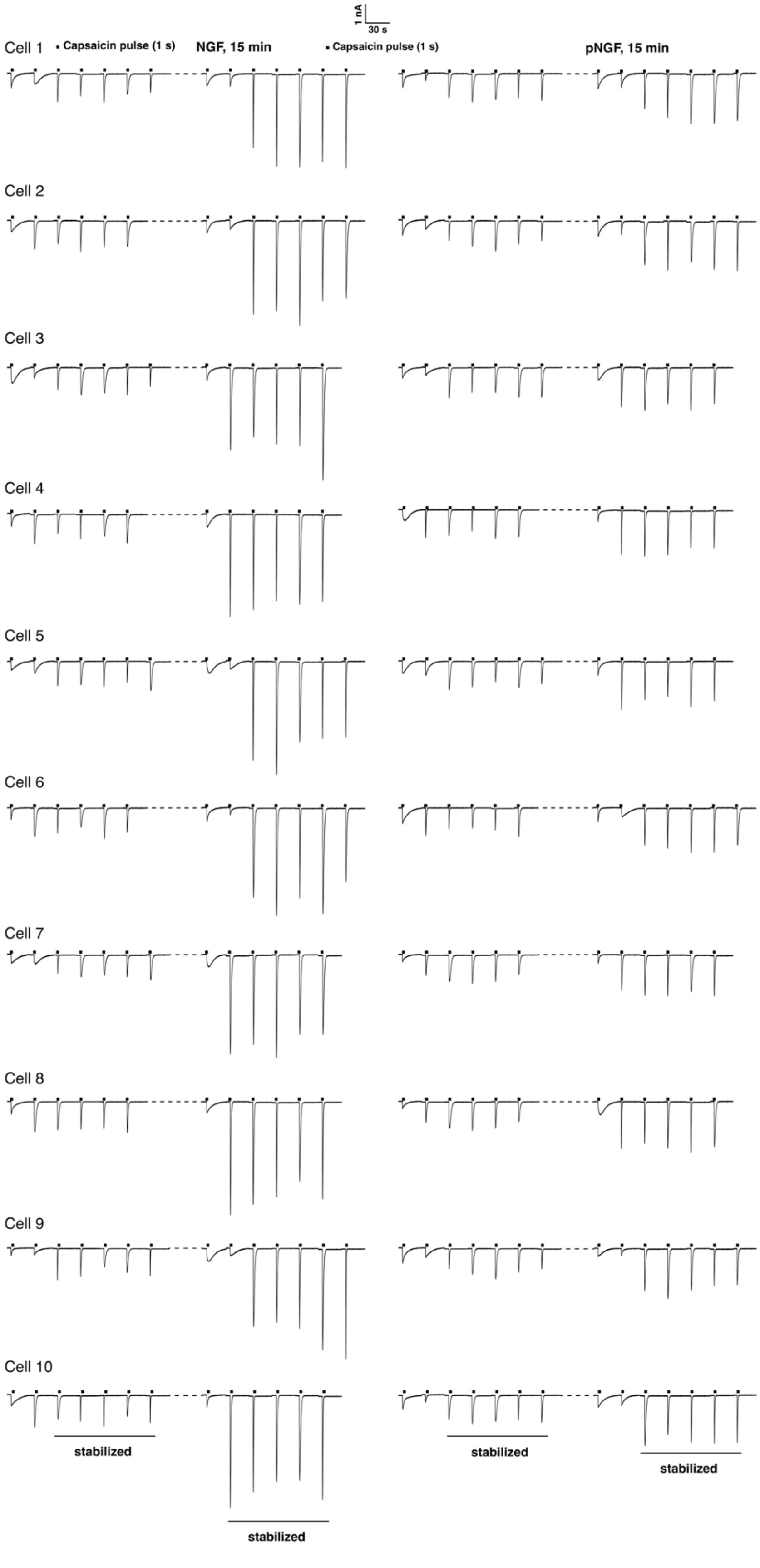


**Supplementary Figure 2**: **Capsaicin-evoked currents in TRPV1 upon treatment with NGF and NGF$^{painless}$**. Representative capsaicin-evoked inward currents (5 times of 1 sec pulses of 10 μM capsaicin at an interval of 30 sec post stabilization, as highlighted) in N = 10 voltage-clamped TrkA- and TRPV1-co-expressing HEK293 cells before and after treatment with 100 ng/ml of NGF (left panels) or NGF$^{painless}$ (right panels) for 15 min at 25°C.

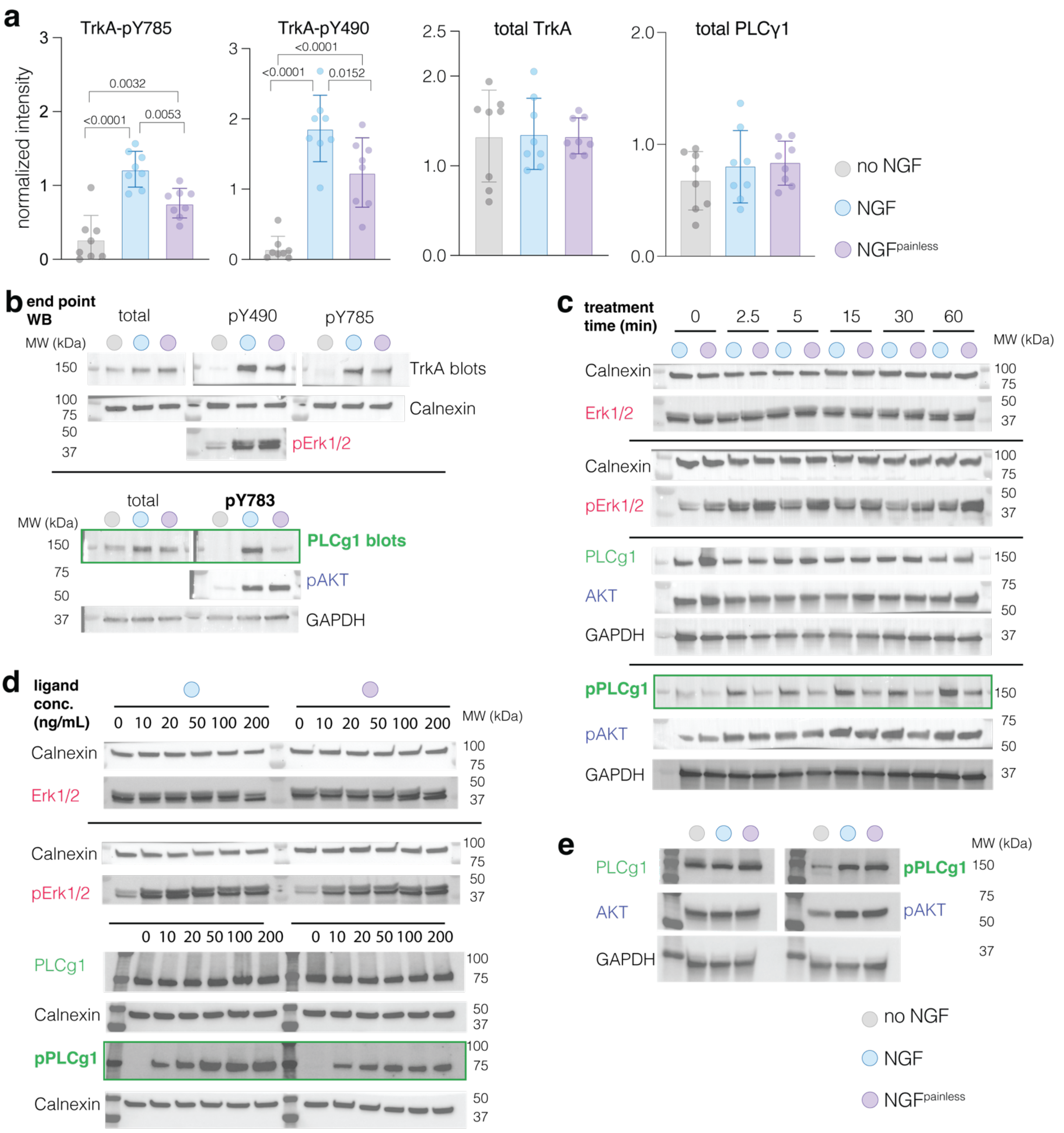


**Supplementary Figure 3**: **NGF- and NGF$^{painless}$-mediated TrkA signaling**. **a.** Western blot analysis comparing levels of TrkA autophosphorylation at Y490 and Y785 and the total levels of TrkA and PLCγ1 following stimulation of TrkA-expressing SHSY5Y cells with NGF or NGF$^{painless}$ for 15 min. No treatment is used as a control. Phosphorylation levels for each protein were quantified using band densitometry and normalized with respect to the loading control using Fiji. Data is shown as mean ± s.d. from N = 8 independent experiments. Results of one-way ANOVA analysis and corresponding P-values are reported. The P-values defined as: *, <0.05; **, <0.1; ***, <0.001; ****, <0.0001. **b.** Representative western blots showing the total levels of TrkA and PLCγ1, as well as the phosphorylation levels in TrkA (pY490 and pY785), PLCγ1 (pY783), pERK, and pAKT using specific antibodies against the total protein or phospho-specific antibodies, respectively. The amount of calnexin or

GAPDH was used as a loading control, depending on the molecular weight of the other species being analyzed. Data is shown for cells under three different conditions: no NGF or cells treated with 50 ng/mL NGF or $NGF^{painless}$ for 15 min. **c.** Representative western blots showing the phosphorylation levels of PLCγ1 (pY783), pERK, and pAKT when treated with NGF or $NGF^{painless}$ for a range of treatment times (0 – 60 min). Immunoblots showing the total levels of these three proteins are also presented. Calnexin or GAPDH were used as loading controls, depending on the molecular weight of the species being analyzed. **d.** Representative western blots showing the phosphorylation levels of PLCγ1 (pY783) and pERK treated with six different NGF or $NGF^{painless}$ concentrations (ranging from 0-200 ng/mL) for 15 min. Immunoblots showing the total levels of these two proteins are also presented. Calnexin or GAPDH were used as loading controls, depending on the molecular weight of the species being analyzed. **e.** Representative western blots showing the phosphorylation levels of PLCγ1 and AKT, along with total levels of these proteins, when cells expressing the $TrkA^{ERDA}$ mutant receptor is treated with 50 ng/ml of either NGF or $NGF^{painless}$ for 15 minutes. GAPDH was used as a loading control.

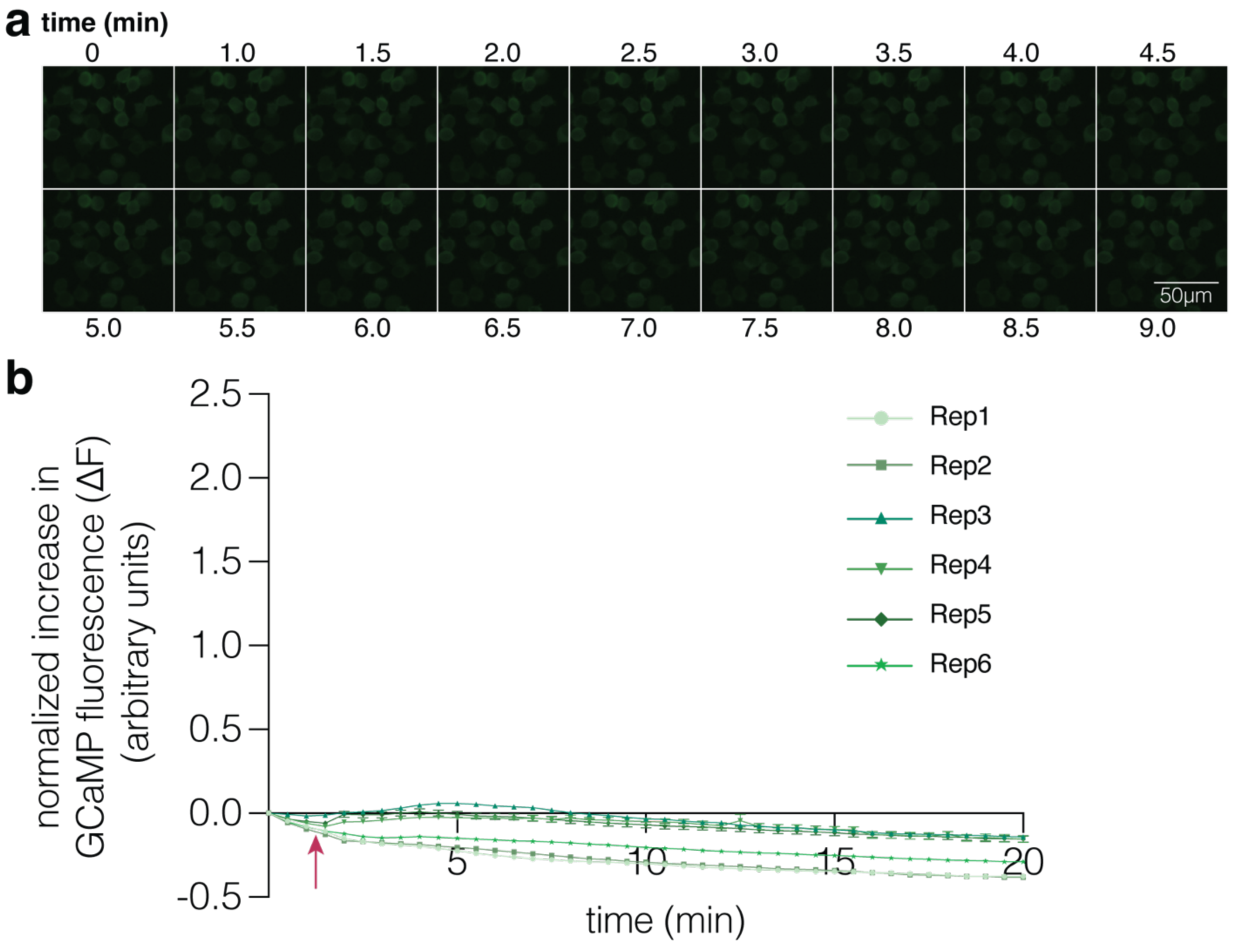


**Supplementary Figure 4**: **Establishing the specificity of the GCaMP8 biosensor**. **a.** Representative images showing GCaMP fluorescence upon treatment of TrkA-expressing SHSY5Y cells with the imaging media (negative control) over 9 minutes. **b.** Plot of GCaMP8 fluorescence change over a period of 20 min showing no change. The red arrow indicates the addition of imaging media. The GCaMP intensity is averaged for 500 - 1500 single cells (shown as mean ± SEM) in each of N = 6 independent experiments.

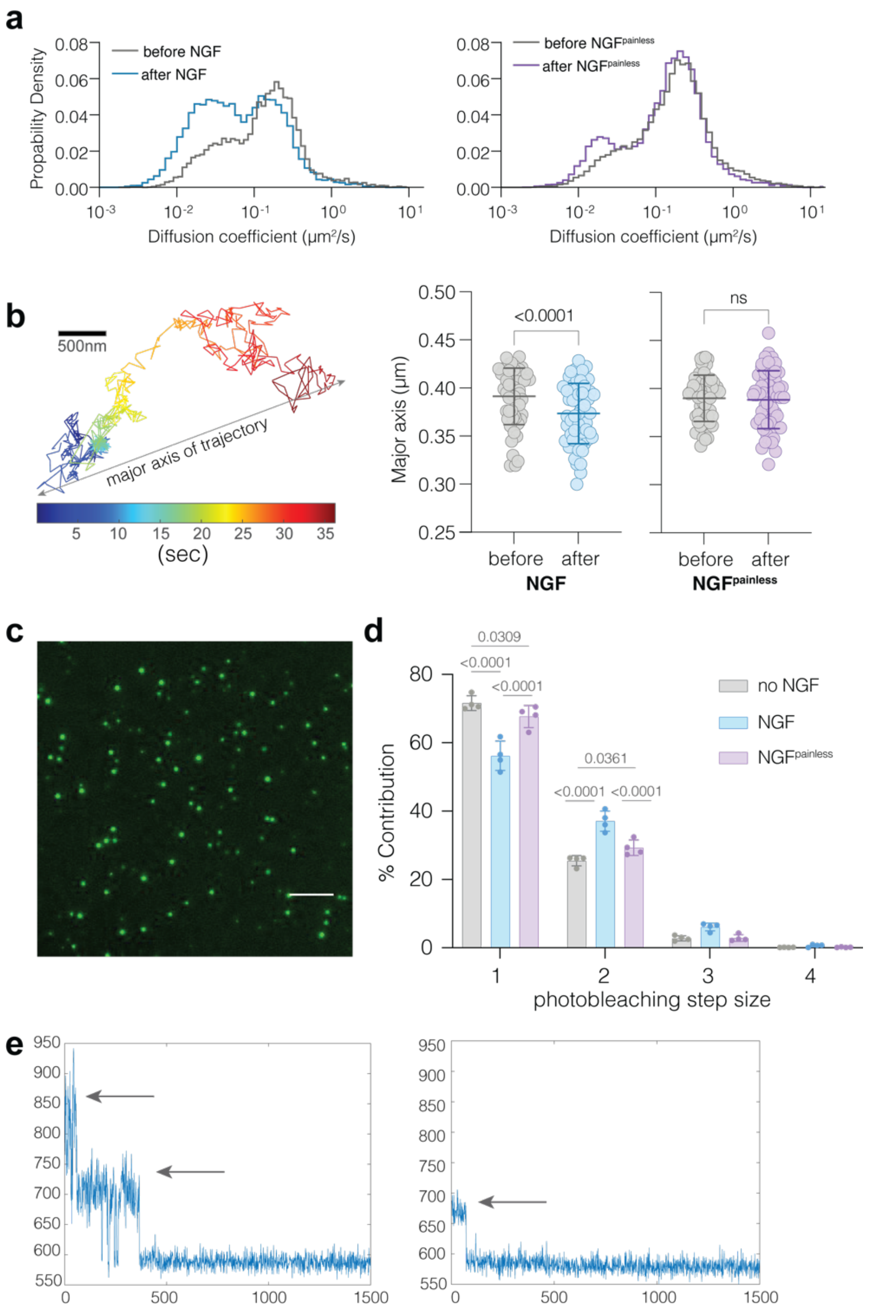


**Supplementary Figure 5**: **Single-Particle Tracking (SPT) and Native-nanoBleach analyses analyses of TrkA upon treatment with NGF or NGF$^{painless}$. a.** Rolling-window diffusion coefficient histograms from a representative single cell are compared before (grey) and after 15-minute incubations with NGF (**left**) or PL-NGF (**right**). **b.** Average length of the major axis of TrkA trajectory ellipses from N = 56 or 54 single cells before (grey circles) and after (blue circles) treatment with NGF (**left**) or PL-NGF (purple circles, **right**), respectively. The major axis is defined as depicted on a representative trajectory (left panel). The data is shown as mean ± s.d. Two-sided paired t-test-derived P values are reported. **c.** Representative single-molecule total internal reflection fluorescence (TIRF) image, where a single green spot represents a single native nanodisc-target protein complex at laser power 17 mW and exposure of 80 msec. Scale bar, 5 μm. **d.** Representative photobleaching traces showing a decrease in GFP intensity over time for proteins bearing two (left) and one (right) mature GFP subunits. The steps in each trace are highlighted using arrows. **e.** Experimentally obtained GFP-photobleaching step distribution of TrkA in native nanodiscs in the presence and absence of either NGF or NGF$^{painless}$. Data is shown from a total of about 3000-4000 single nanodiscs from N = 4 biologically independent samples and represented as mean ± s.d. Results of two-way ANOVA analysis showing differences in step size distributions with P-values are reported.

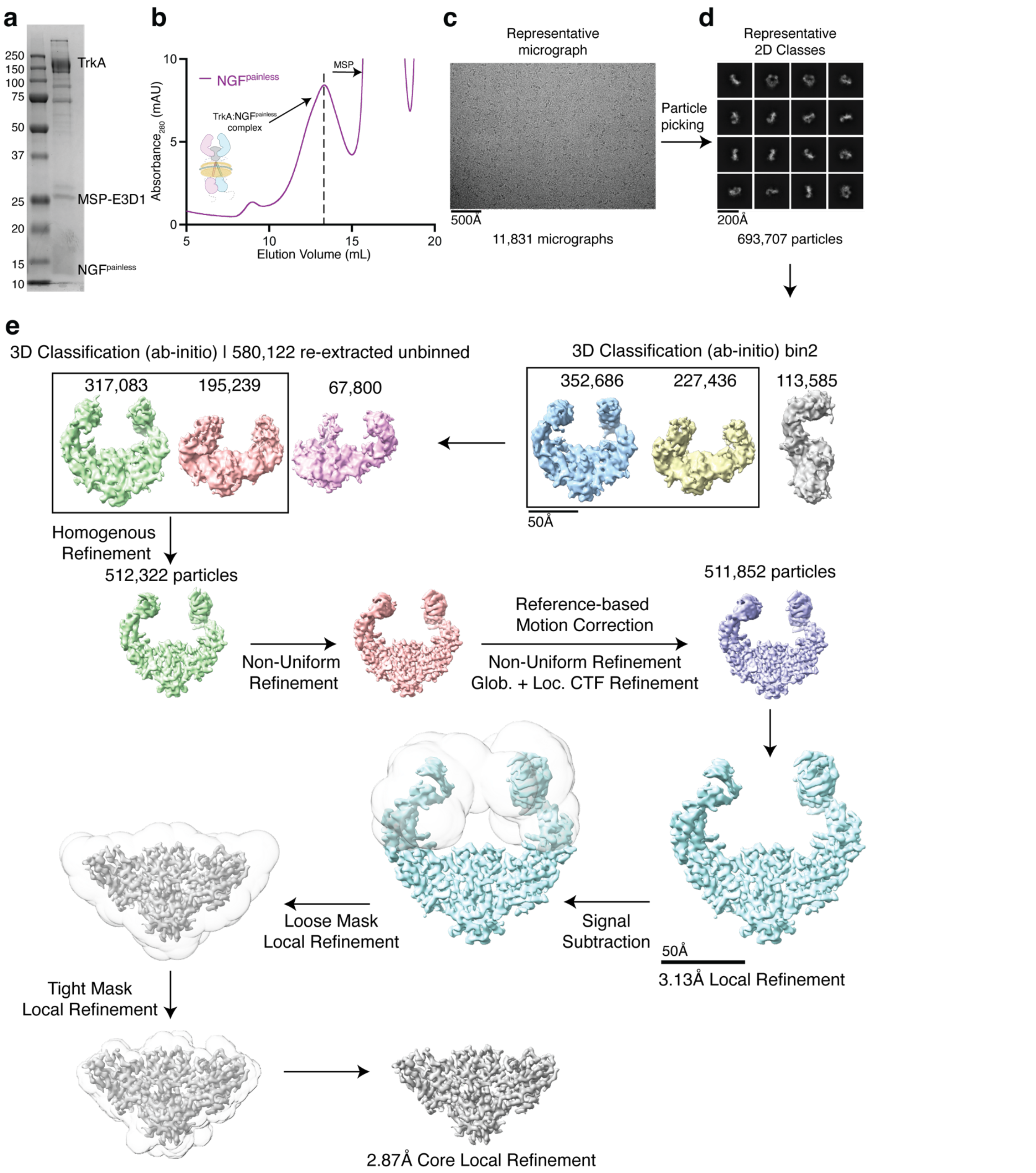


**Supplementary Figure 6**: **Single particle cryo-electron microscopy (cryo-EM) workflow for 2:2 TrkA:NGF$^{painless}$ complex. a.** SDS-PAGE (4-20%) analysis of the nanodisc reconstituted TrkA:NGF$^{painless}$ sample showing distinct bands for TrkA, MSP-E3D1, and NGF$^{painless}$. **b**. Size exclusion chromatography trace of the TrkA:NGF$^{painless}$**,** showing the peak at 13.32 ml (Superose 6 Increase 10/300 GL column), which was used for cryo-EM analysis. **c**. A representative motion-corrected micrograph showing the TrkA:NGF$^{painless}$ complexes in vitreous ice. Scale bar, 500Å**.** The dataset was entirely processed in cryoSPARC v.4.6.2. **d**. Representative 2D class averages showing various orientations of the TrkA:NGF$^{painless}$ complex**.** 4.28 million particles were selected using the template picker, cleaned by 2D classification to remove obvious junk particles, and then pooled. **e.** Duplicates were removed, and the particles were sorted by iterative 2D and 3D classification. Throughout the processing workflow, classes with the most well-resolved extracellular domain (ECD) were used as input references for the subsequent steps of processing. A final clean stack of 511,852 particles was used for downstream 3D analysis. C1 symmetry was applied throughout data processing to avoid artefacts from symmetrization and to capture the dynamics of the ECD arms of TrkA. TrkA ECD-Ig2 density was signal subtracted from the final particle set and refined further to obtain a structure of core (TrkA:NGF$^{painless}$ interface) at 2.87 Å.

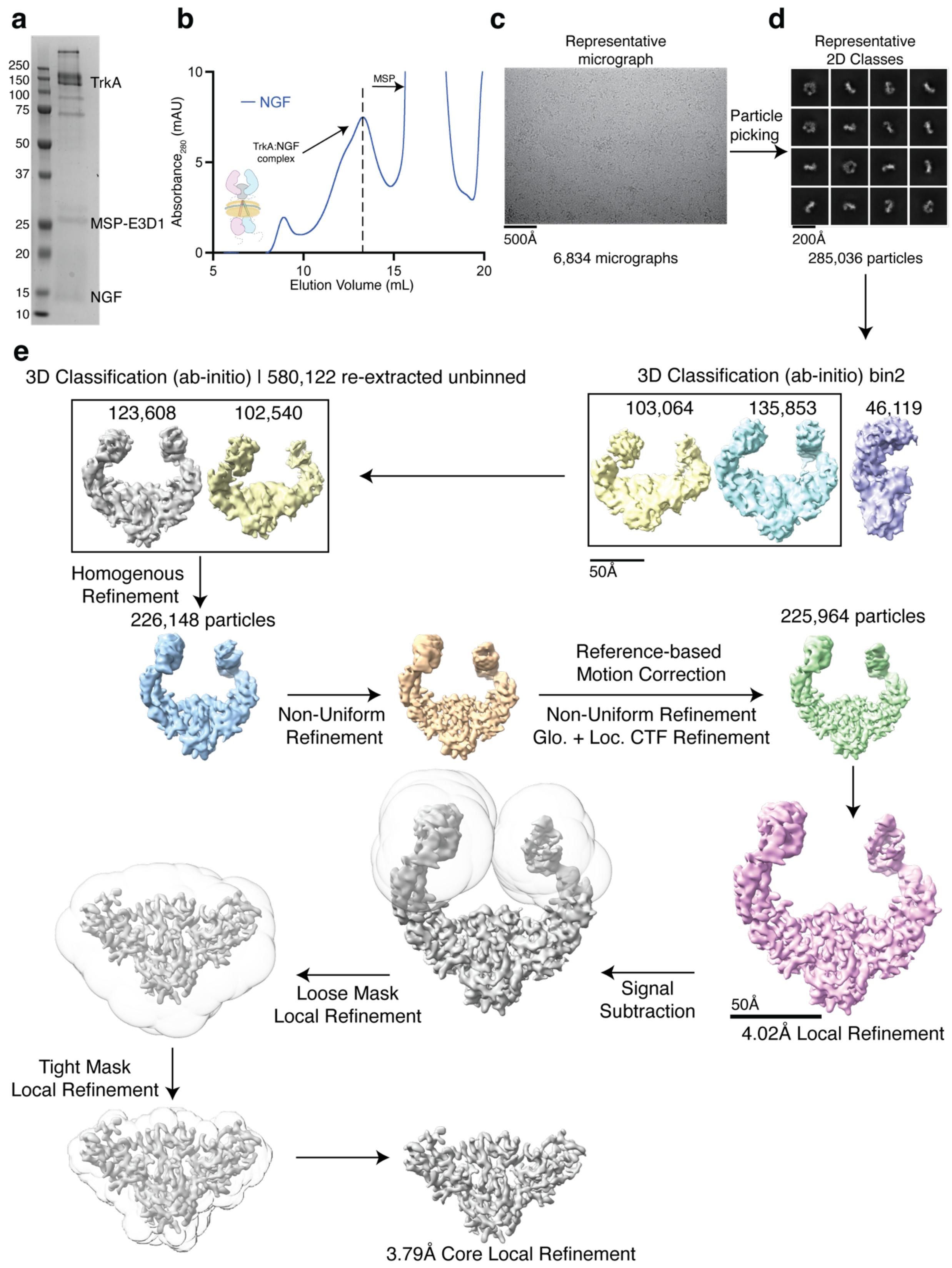


**Supplementary Figure 7**: **Single particle cryo-electron microscopy (cryo-EM) workflow for 2:2 TrkA:NGF complex. a.** SDS-PAGE (4-20%) analysis of the nanodisc reconstituted TrkA:NGF sample showing distinct bands for TrkA, MSP-E3D1, and NGF. **b**. Size exclusion chromatography trace of the TrkA:NGF**,** showing the peak at 13.30 ml (Superose 6 Increase 10/300 GL column), which was used for cryo-EM analysis. **c**. A representative motion-corrected micrograph showing the TrkA:NGF complexes in vitreous ice. Scale bar, 500Å**.**

The dataset was entirely processed in cryoSPARC v.4.6.2. **d**. Representative 2D class averages showing various orientations of the TrkA:NGF complex**.** 2.43 million particles were selected using the template picker, cleaned by 2D classification to remove obvious junk particles, and then pooled. **e.** Duplicates were removed, and the particles were sorted by iterative 2D and 3D classification. Throughout the processing workflow, classes with the most well-resolved extracellular domain (ECD) were used as input references for the subsequent steps of processing. A final clean particle stack of 225,964 particles was used for downstream 3D analysis. C1 symmetry was applied throughout data processing to avoid artefacts from symmetrization and to capture the dynamics of the ECD arms of TrkA. TrkA ECD-Ig2 density was signal subtracted from the final particle set and refined further to obtain a structure of core (TrkA:NGF interface) at 3.79 Å.

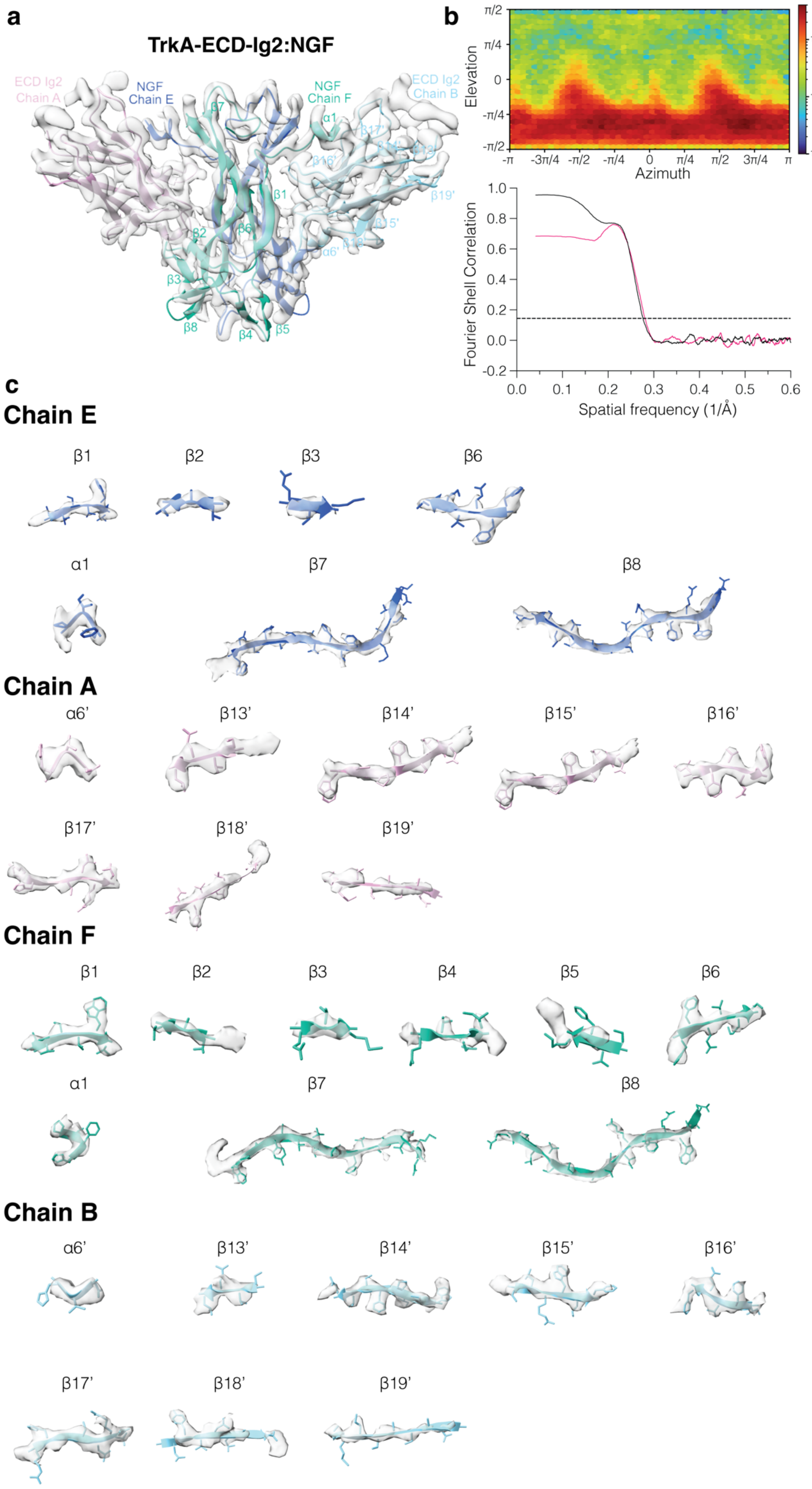
a
TrkA-ECD-Ig2:NGF
ECD Ig2 Chain A
NGF Chain E
NGF Chain F
ECD Ig2 Chain B
β7
α1
β17'
β16'
β19'
β1
β15'
β2
β6
α6'
β3
β8
β4
β5
b
π/2
π/4
0
-π/4
-π/2
Elevation
-π
-3π/4
-π/2
-π/4
0
π/4
π/2
3π/4
π
Azimuth
### of images
10²
10¹
10⁰
1.0
0.8
0.6
0.4
0.2
0.0
-0.2
Fourier Shell Correlation
0.0
0.1
0.2
0.3
0.4
0.5
0.6
Spatial frequency (1/Å)
c
Chain E
β1
β2
β3
β6
α1
β7
β8
Chain A
α6'
β13'
β14'
β15'
β16'
β17'
β18'
β19'
Chain F
β1
β2
β3
β4
β5
β6
α1
β7
β8
Chain B
α6'
β13'
β14'
β15'
β16'
β17'
β18'
β19'

**Supplementary Figure 8**: **Cryo-EM data quality, reconstruction, and model building for TrkA-ECD-Ig2:NGF. a**. The fit of the deposited model (PDB: 9YKU), showing TrkA ECD Ig2 chains in light blue and pink and the NGF chains in dark blue and green, into the refined map (EMD-73058, gray). All maps are visualized in ChimeraX (version 1.7) and displayed at an isosurface threshold level of 0.08. **b**. Gold standard Fourier shell coefficient (FSC) curve (black) and map to model FSC (pink) are shown. **c.** Cryo-EM density of the secondary structure elements, β1-3,6-8 and α1 for chain E and β1-8 and α1 for chain F, and β13'-19' and α6' for chains A and B, are colored per chain. Side chains for almost all the elements could be modeled unambiguously into the density. At places with missing density, the side chains were trimmed to Cβ.

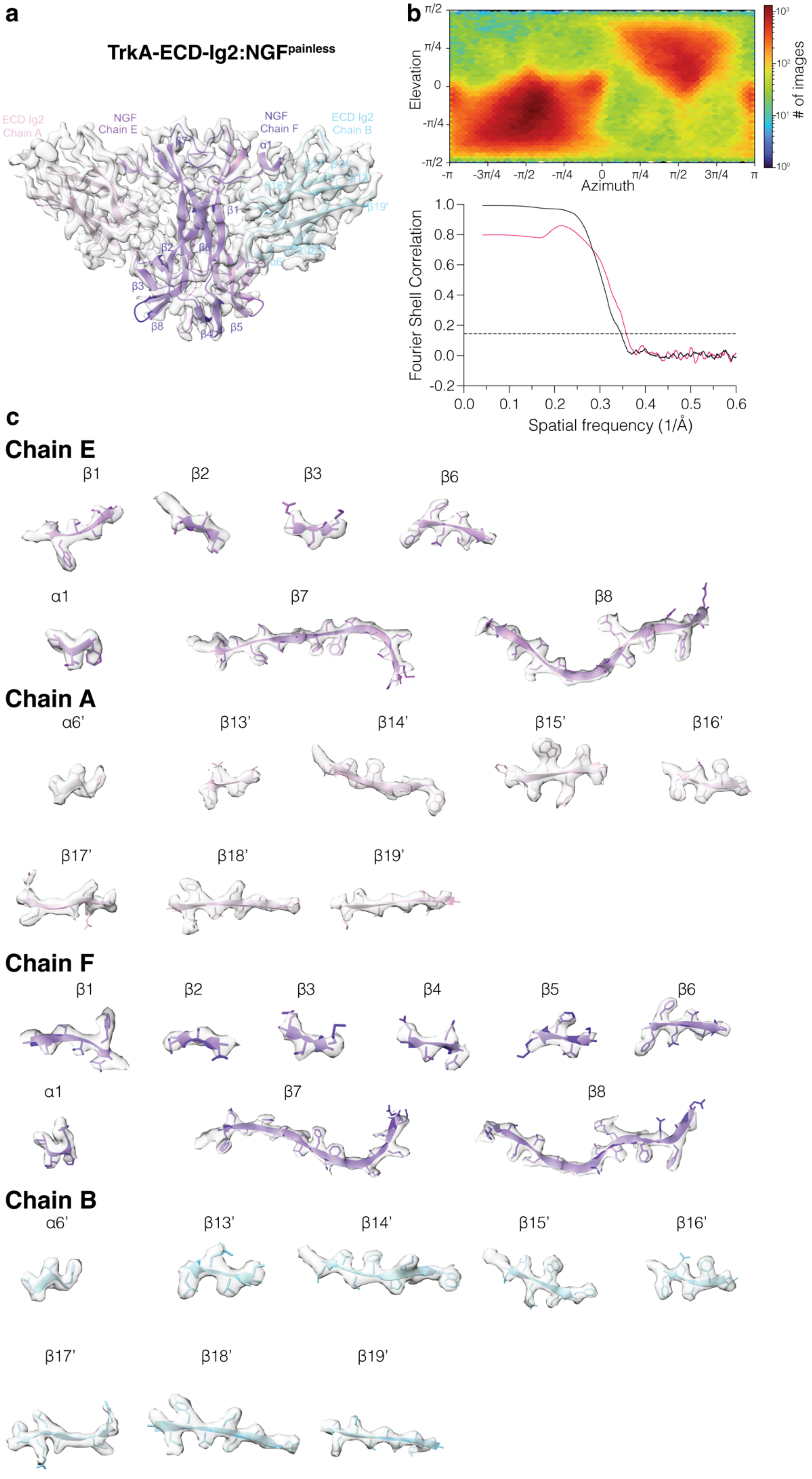

a
TrkA-ECD-Ig2:NGFpainless
ECD Ig2 Chain A
NGF Chain E
NGF Chain F
ECD Ig2 Chain B
b
Elevation
Azimuth
### of images
Fourier Shell Correlation
Spatial frequency (1/Å)
c
Chain E
β1
β2
β3
β6
α1
β7
β8
Chain A
α6'
β13'
β14'
β15'
β16'
β17'
β18'
β19'
Chain F
β1
β2
β3
β4
β5
β6
α1
β7
β8
Chain B
α6'
β13'
β14'
β15'
β16'
β17'
β18'
β19'

**Supplementary Figure 9**: **Cryo-EM data quality, reconstruction, and model building for TrkA-ECD-Ig2:NGF$^{painless}$.** **a**. The fit of the deposited model (PDB: 9YKT), showing TrkA ECD Ig2 chains in light blue and pink and the NGF$^{painless}$ chains in violet and purple, into the refined map (EMD-73057, gray). Maps are visualized in ChimeraX (version 1.7) and displayed at an isosurface threshold level of 0.08. **b**. Gold standard Fourier shell coefficient (FSC) curve (black) and map to model FSC (pink) are shown. **c.** Cryo-EM density of the secondary structure elements, β1-3, 6-8 and α1 for chain E and β1-8 and α1 for chain F, and β13'-19' and α6' for chains A and B, are colored per chain. Side chains for almost all the elements could be modeled unambiguously into the density. At places with missing density, the side chains were trimmed to Cβ.

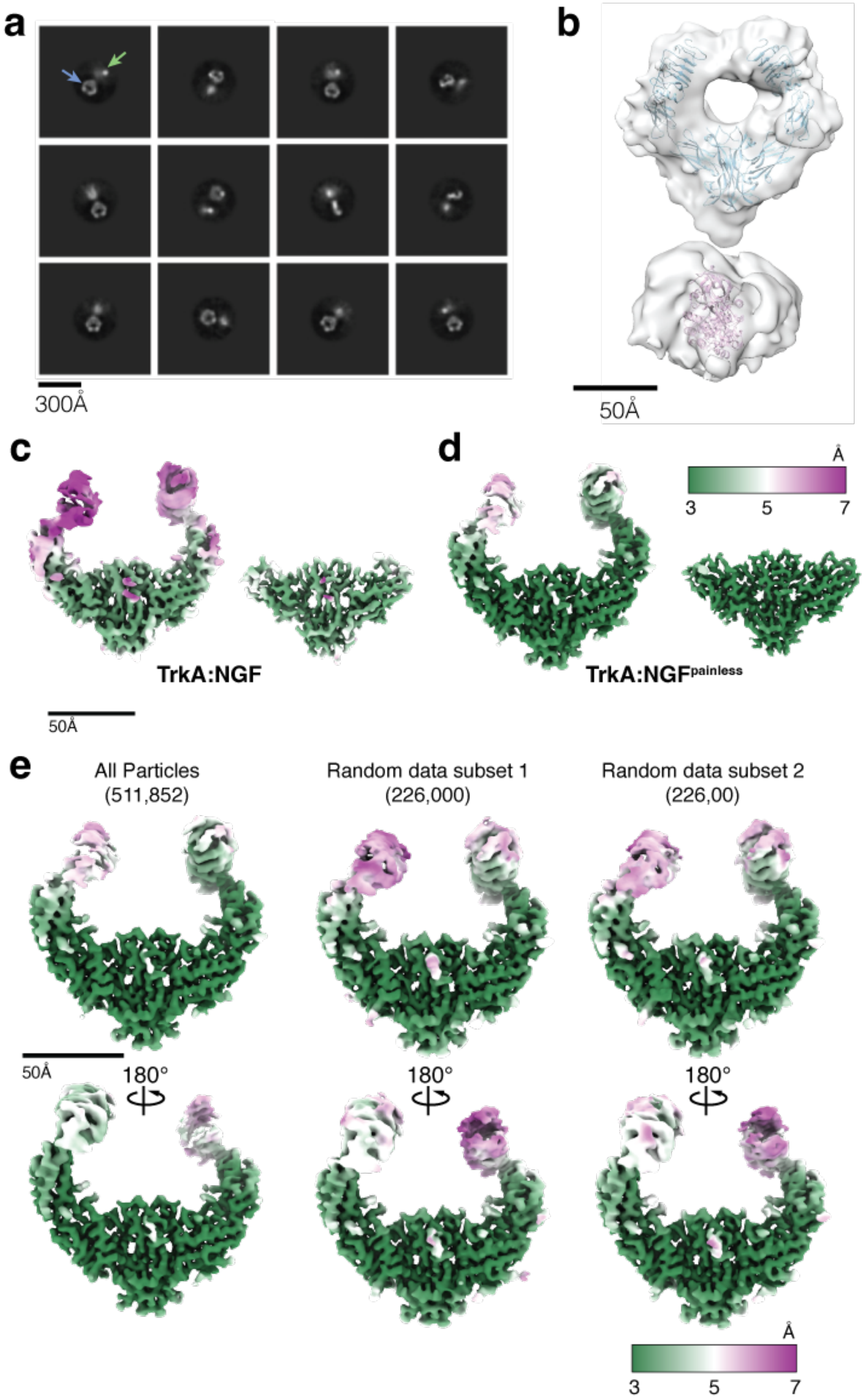


**Supplementary Figure 10**: **Structural analysis of full-length TrkA:NGF complex. a.** Representative 2D class averages of the full-length TrkA:NGF complex showing density for the extracellular domain (ECD, blue arrow) and the kinase domain (KD, green arrow). Owing to flexibility, the kinase domain is poorly resolved. Scale bar, 300 Å. **b.** Low-resolution 3D reconstruction of the full TrkA:NGF complex with rigid body fit of PDB:2IFG (truncated ECD:NGF crystal structure), 4F0I (truncated KD crystal structure). Scale bar, 50 Å. **c.** Comparison of local resolution maps for TrkA:NGF and TrkA:NGF$^{painless}$. Left: Local resolution map of the TrkA:NGF complex. Inset shows local resolution map of the NGF+ECD-Ig2 of TrkA core region formed by the receptor-ligand interface. Scale bar, 50 Å. **d.** Left: Local resolution map of the TrkA:NGF$^{painless}$ complex. Inset shows local resolution map of the NGF$^{painless}$+ECD-Ig2 of TrkA core region formed by the receptor-ligand interface. Scale bar, 50 Å. **e.** Views of the local resolution maps for all particles of the TrkA:NGF$^{painless}$ complex and those from the independent analysis of two randomly selected subsets of these data, with approximately half the number of particles in each case. Scale bar, 50 Å and a colormap are shown.
