## Supplementary Tables for "A painless nerve growth factor variant uncouples nociceptive and neurotrophic TrkA signaling"

**Supplementary Table 1: Cryo-EM data collection, refinement and validation statistics**

| | TrkA:NGF$^{painless}$ (EMD-73057) (PDB 9YKT) | TrkA:NGF (EMD-73058) (PDB 9YKU) |
|---|---|---|
| **Data collection and processing** | | |
| Magnification | 105000x | 105000x |
| Voltage (kV) | 300 | 300 |
| Electron exposure (e-/Å$^2$) | 50 | 50 |
| Defocus range (μm) | 1.0-2.5 | 1.0-2.5 |
| Pixel size (Å) | 0.832 | 0.832 |
| Symmetry imposed | C1 | C1 |
| Initial particle images (no.) | 1081972 | 489958 |
| Final particle images (no.) | 511852 | 225964 |
| Map resolution (Å) | 2.87 | 3.79 |
| FSC threshold | 0.143 | 0.143 |
| **Refinement** | | |
| Initial model used (PDB code) | 2IFG | 2IFG |
| Model resolution (Å) | 3.15 | 4.28 |
| FSC threshold | 0.5 | 0.5 |
| Map sharpening B factor (Å$^2$) | -80 | -80 |
| Model composition | | |
| Nonhydrogen atoms | 3156 | 3160 |
| Protein residues | 402 | 402 |
| B factors (Å$^2$) | | |
| Protein (min/max/mean) | 21.17/120.04/53.80 | 11.88/43.65/23.55 |
| R.m.s. deviations | | |
| Bond lengths (Å) | 0.006 | 0.007 |
| Bond angles (°) | 0.932 | 1.15 |
| Validation | | |
| MolProbity score | 1.72 | 2.2 |
| Clashscore | 5.67 | 11.64 |
| Poor rotamers (%) | 0.29 | 0.58 |
| Ramachandran plot | | |
| Favored (%) | 93.85 | 87.18 |
| Allowed (%) | 6.15 | 12.82 |
| Disallowed (%) | 0 | 0 |

**Supplementary Table 2: Plasmid list (monomeric eGFP is denoted as GFP throughout)**

| Plasmid name | Tag + Position of tag | Uniprot/Addgene_ID/ VectorBuilder ID | Plasmid reference |
|---|---|---|---|
| pEG BacMam | Cterm 3C-GFP-8xHis | Addgene_160687 | PMID: 25299155 |
| pEG BacMam TrkA-3C-GFP-8xHis | Cterm 3C-GFP-8xHis | P04629 | PMID: 38012273 |
| pEG BacMam TrkA-3C-GFP-3xFLAG | Cterm 3C-GFP-3xFLAG | P04629 | Current study |
| pEG BacMam ALFA-linker-TrkA-3C-GFP-3xFLAG | Nterm ALFA-linker Cterm 3C-GFP-3xFLAG | | Current study |
| pSA75-pHRSIN-TrkA-3C-GFP-3xFLAG | Cterm 3C-GFP-3xFLAG | pSA75-pHRSIN backbone is a gift from Dr. Shalini Low-Nam, Purdue University | Current study |
| pSA75-pHRSIN-TrkA-3C-GFP-8xHis | Cterm 3C-GFP-8xHis | | Current study |
| pSA75-pHRSIN-ALFA-linker-TrkA-3C-GFP-3xFLAG | Nterm ALFA-linker Cterm 3C-GFP-3xFLAG | | Current study |
| pLV[Exp]-Puro-EF1A jGCaMP8m | | VectorBuilder ID: VB900169-8014wfn | Current study |
| pLV[Exp]-mCherry/Puro-EF1A>hTRPV1 | | VectorBuilder ID: VB900181-7765gub | Current study |
| pEXQV | | Gift from Dr. Shalini Low-Nam, Purdue University | |
| pMD2.G | | Addgene ID: 12259 | |
| pMDLg/pRRE | | Addgene ID: 12251 | |
| pRSV-Rev | | Addgene ID: 12253 | |
| pGP-CMV-jGCaMP8m | | Addgene ID: 162372 | |

**Supplementary Table 3: List of cell lines**

| Cell type | Gene integrated | Source |
|---|---|---|
| SHSY5Y | | ATCC-CRL-2266 |
| SHSY5Y-TrkA | TrkA | Kerafast-ECP006 |
| HEK293FT | NA | Gift from Dr. John Kuriyan's lab |
| Expi293 | NA | Gift from Dr. Karin Reinisch's lab |
| Sf9 insect cells | NA | ATCC-CRL1711 |
| Sf9 Easy Titer cells | NA | ATCC-CRL-3357 |
| HEK293FT-TrkA-3C-GFP-8xHis | TrkA-3C-GFP-8xHis | current study |
| SHSY5Y-TrkA-3C-GFP-8xHis | TrkA-3C-GFP-8xHis | current study |
| SHSY5Y-ALFA-linker-TrkA-3C-GFP-3xFLAG | ALFA-linker-TrkA-3C-GFP-3xFLAG | current study |
| HEK293FT-TrkA-GFP-8xHis + hTRPV1 + mCherry | TrkA-GFP-8XHis + hTRPV1 + mCherry | current study |
| SHSY5Y-TrkA-jGCaMP8m | jGCaMP8m | current study |

**Supplementary Table 4: Bacterial strains**

| Cell type | Company | Catalog number |
|---|---|---|
| *E. coli* -TOP10 competent cells | ThermoFisher Scientific | C404010 |
| Max Efficiency DH10Bac Competent Cells | ThermoFisher Scientific | 10361012 |
| NEB Stable Competent *E. coli* (High Efficiency | NEB | C3040H |
| BL21 (DE3) Competent Cells | ThermoFisher Scientific | EC0114 |

**Supplementary Table 5: List of antibodies**

| Antibody | Species/type | Company | Catalog Number |
|---|---|---|---|
| TrkA Antibody | Rabbit/polyclonal | Cell Signaling Technology | 2505S |
| Phospho-TrkA (Tyr490)/TrkB (Tyr516) (C35G9) | Rabbit/monoclonal | Cell Signaling Technology | 4619S |
| Phospho-TrkA (Tyr785)/TrkB (Tyr816) (C67C8) | Rabbit/monoclonal | Cell Signaling Technology | 4168S |
| Phospho-TrkA (Tyr674/675) | Rabbit/monoclonal | Cell Signaling Technology | 4621S |
| p44/42 MAPK (Erk1/2) (137F5) | Rabbit/monoclonal | Cell Signaling Technology | 4695S |
| Phospho-p44/42 MAPK (Erk1/2) (Thr202/Tyr204) (D13.14.4E) XP | Rabbit/monoclonal | Cell Signaling Technology | 4370 |

| | | | |
|---|---|---|---|
| Akt (pan) (C67E7) | Rabbit/monoclonal | Cell Signaling Technology | 4691S |
| Phospho-Akt (Ser473) (D9E) XP | Rabbit/monoclonal | Cell Signaling Technology | 4060S |
| Calnexin | Rabbit/monoclonal | Abcam | ab92573 |
| GAPDH (14C10) | Rabbit/monoclonal | Cell Signaling Technology | 2118S |
| PLCγ1 | Rabbit/polyclonal | Cell Signaling Technology | 2822S |
| Phospho-PLCγ1 (Tyr783) | Rabbit/polyclonal | Cell Signaling Technology | 2821S |
| FluoTag®-X2 anti-ALFA, Alexa Fluor 647 | | NanoTag Biotechnologies | N1502-AF647-L |
| GFP VHH, biotinylated recombinant binding protein | | ChromoTek | gtb-250 |

**Supplementary Table 6: Chemicals & Reagents**

| **Reagent** | **Company** | **Catalog Number** |
|---|---|---|
| Trizma base | Sigma-Aldrich | T6066-1KG |
| 10x Tris/Glycine/SDS Electrophoresis Buffer | Bio-Rad | 1610732EDU |
| Sodium Chloride | Sigma-Aldrich | S9888-500G |
| Potassium Chloride | Sigma-Aldrich | 746436-500g |
| Glycerol | Sigma-Aldrich | G7893-1L |
| Phenylmethylsulfonyl fluoride (PMSF) | Sigma-Aldrich | 52332-5GM |
| Pepstatin A | AmericanBio | ab01555-00010 |
| Sodium Fluoride | Sigma-Aldrich | 201154-100G |
| Sodium Orthovanadate | EMD Millipore | 567540-5GM |
| Sodium Butyrate | Sigma Aldrich, | 303410 |
| Methanol | Sigma-Aldrich | 179337-500ML |
| Soy phospholipid mixture | Sigma-Aldrich | 11145-50G |
| Decyl Maltose Neopentyl Glycol | Anatrace | NG322-5-GM |
| Ethanol | Sigma Aldrich | E7023-500ML |
| 3X FLAG peptide | Elim biopharma | 24200 |
| SMALP 200 | Aurorium | SMALP200 |
| Clarity Western ECL substrate | Bio-Rad | 1705000 |
| Triton X-100 | Sigma-Aldrich | T8787-250ML |
| Bovine Serum albumin | Sigma-Aldrich | A9647-100G |
| Chloroform | Sigma-Aldrich | |
| Chloroform | Sigma-Aldrich | 319988 |
| Isopropanol | Sigma-Aldrich | 190764 |
| Poly-L-Lysine | EMD Millipore | A-005-C |

| | | |
|---|---|---|
| Polyethylenimine (PEI) | Sigma-Aldrich | 764965 |
| PLL-PEG/PEG-biotin | SuSoS | PLL(20)-g[3.5]-PEG(2) |
| Streptavidin | NEB | N7021S |
| Capsaicin | Sigma-Aldrich | M2028-50MG |
| HEPES | Sigma-Aldrich | H3375-1KG |
| Calcium chloride | Sigma-Aldrich | 223506-500G |
| D-mannitol | Sigma-Aldrich | M4125-100G |
| Potassium fluoride | Spectrum Chemical MFG Corp. | P1621-500G |
| EGTA | Fluka | 03779-10G |
| Na2ATP | Sigma-Aldrich | A2383-1G |
| $Li_2GTP$ | Sigma-Aldrich | G5884-100MG |
| Potassium hydroxide | Sigma-Aldrich | 221473-500G |
| Ammonium acetate | MP Biomedical | 198759 |
| Decyl Maltose Neopentyl Glycol (DMNG) | Anatrace | NG322 |

**Supplementary Table 7: Media & Buffers**

| Reagent | Company | Catalog Number |
|---|---|---|
| Expi293 Expression Medium | Thermo Fisher Scientific | A1435101 |
| DMEM | Thermo Fisher Scientific | 11965092 |
| Advanced DMEM/F12 medium | Thermo Fisher Scientific | 12634010 |
| Opti-MEM Reduced Serum Medium | Thermo Fisher Scientific | 31985070 |
| Sf-900 II SFM media | Thermo Fisher Scientific | 10902088 |
| Antibiotic-Antimycotic | Thermo Fisher Scientific | 15240062 |
| DPBS | Thermo Fisher Scientific | 14190144 |
| Trypsin (0.05%) | Thermo Fisher Scientific | 25300-054 |
| Fetal bovine serum | Sigma-Aldrich | 12306C-500ML |
| L-Glutamine | Thermo Fisher Scientific | 25030081 |
| Detachin™ cell detachment solution | Avantor | MSPP-T100110 |
| Typhane Blue stain (0.4%) | Gibco | 15250-061 |
| Polybrene | EMD Millipore | TR-1003-G |
| DMEM/F12, no phenol red | Thermo Fisher Scientific | 21041-025 |
| RPMI 1640 Medium | Thermo Fisher Scientific | 11875-093 |
| RPMI 1640 Medium, no phenol red | Thermo Fisher Scientific | 11835-030 |

**Supplementary Table 8: Inhibitors and Antibiotics**

| Reagent | Company | Catalog Number |
|---|---|---|

| | | |
|---|---|---|
| Protease Inhibitor | Thermo Fisher Scientific | A32953 |
| cOmplete Protease Inhibitor | Sigma-Aldrich | 11836170001 |
| TrkA inhibitor (GW441756) | Selleck Chemicals | S2891 |
| Ampicillin | GoldBio | A-301-25 |
| Kanamycin | GoldBio | K120-50 |
| Tetracycline Hydrochloride | Sigma-Aldrich | T7660-5G |
| Gentamicin | Gibco | 15710-064 |
| Geneticin | Gibco | 10131-035 |

### Supplementary Table 9: Beads & Resins

| Reagent | Company | Catalog Number |
|---|---|---|
| ANTI-FLAG M2 Affinity Gel | Sigma-Aldrich | A2220-5ML |
| Bio-Beads SM-2 Adsorbents | Bio-Rad | 1528920 |

### Supplementary Table 10: Miscellaneous items

| Items | Company | Catalog Number |
|---|---|---|
| Spin-X centrifuge tube filter | Corning | 8160 |
| Vivaspin 20, 100 kDa | Cytiva | 28932363 |
| Vivaspin 6. 100 kDa | Cytiva | 28932319 |
| Ultra Au Foil R1.2/1.3 300 mesh | EMS | Q350AR13A |
| 4–20% Mini-PROTEAN® TGX Stain-Free™ Protein Gels, | Bio-Rad | 4568096 |
| Trans-Blot turbo RTA transfer kit, PVDF | Bio-Rad | 1704272 |
| PVDF membrane | Bio-Rad | 1620177 |
| Ibidi Sticky-Slide | Ibidi | 80608 |
| Coverslips | Ibidi | 10812 |
| MatTek Dishes | MATTEK | P35GF-1.5-14-C |
| Zeba spin desalting columns | Thermo Fisher Scientific | 89877 |
| 8-16% Mini-PROTEAN® TGX Stain-Free™ Protein Gels, | Bio-Rad | 4561106 |
| IPTG | GoldBio | 12481C25 |

### Supplementary Table 11: List of buffers for membrane protein purification

| Buffer type | Buffer composition |
|---|---|
| *Lysis buffer | 25 mM Tris-HCl pH 8.0, 150 mM KCl |
| [*,$]Membrane resuspension buffer | 20 mM Tris-HCl, pH 8.0, 150 mM NaCl, 5 % glycerol |
| FSEC buffer | 20 mM Tris pH 8.0, 150 mM NaCl |
| **TrkA solubilization buffer using detergent** | |

| | |
|---|---|
| Solubilization buffer | Membrane resuspension buffer, 1% DMNG |
| Wash buffer | Membrane resuspension buffer, 0.1% DMNG |
| Elution buffer | Wash buffer supplemented with 0.2mg/ml FLAG peptide. |

*All lysis and membrane resuspension buffers were supplemented with protease inhibitor cocktail tablets (Pierce, Thermo), PMSF, and phosphatase inhibitors (sodium orthovanadate and sodium fluoride).

[$] For membrane resuspension buffers, 2% of final concentration of copolymers SMA was added.
